# The contribution of epigenetic variation to evolution in crows

**DOI:** 10.1101/2024.05.22.595340

**Authors:** Justin Merondun, Jochen B. W. Wolf

## Abstract

Chromatin modifications provide a substrate for epigenetic variation with evolutionary potential. To quantify the contribution of this layer of variation to evolution we leveraged genome and methylome sequencing data from an incipient avian species: all-black carrion crows, grey-coated hooded crows and their hybrids. Combining controlled experimentation under common garden conditions and sampling of natural genetic variation across the hybrid zone we show that 5mC methylation variation was almost exclusively explained by genome properties and ontogenetic program of the organism. Evidence for an environmental contribution was minor, and all methylation variation of potential importance to speciation clustered in intergenic space within a genomic region of elevated genetic differentiation encoding the diagnostic color-contrast between taxa. We conclude that methylation variation may aid in phenotypic translation of genetic polymorphism, but provides little scope for an autonomous contribution to evolution in this system.

## Introduction

Mutations are the ultimate source of evolution. Prevailing evolutionary theory conceptualizes mutations as random changes to the DNA backbone filtered by selection, depleted by genetic drift, reorganized by recombination and redistributed by migration [1]. The concept of epigenetic inheritance challenges this paradigm, as it introduces a second layer of potentially heritable variation that is not subject to alterations of the nucleotide sequence [2]. DNA methylation is a prime candidate of a molecular epigenetic inheritance system featuring variation along the genome, within and among individuals and populations [3,4]. The underpinnings of variation in DNA methylation are diverse and include i) spontaneous epimutations, ii) environmental induction, iii) physiological processes establishing somatic cell fate [ontogenetic program sensu 5] and iv) genetic constraints imposed by genome properties (chromosomal or genomic features) or the organism’s genotype [6]. While the latter couple of sources are readily incorporated into traditional evolutionary theory, the first two are not. To judge the autonomous potential of DNA methylation for evolution, it is therefore crucial to obtain a quantitative understanding of the sources shaping natural variation [7].

In plants, evolutionarily relevant methylation variation seems rather widespread [8]. There is evidence for spontaneous, random epimutations in DNA methylation [9–11], environmentally-induced epimutations [12] and transgenerational inheritance of both [13–15]. In animals, with strict soma-germline separation and epigenetic reprogramming [16], variation in 5mC methylation that is independent of the genotype is expected to be more limited [17]. While animal research aiming to separate genetic, developmental, and environmental impacts on variation in DNA methylation is gaining traction [18], the evidence for stable inheritance of autonomous chromatin modifications remains scarce [18–21]. Numerous vertebrate studies have linked 5mC methylation variation with environmental change [22–24] and/or phenotypic variation [25], but few have done so while controlling for confounding genetic effects [26–28](see also [29] for transposable elements). In birds, DNA methylation has been associated with stress resilience in tree swallows [30] and urbanization in great tits [31], while conversely a recent well-designed study employing partial cross-fostering under controlled conditions suggests a very limited role for environmental induction on methylation variation independent of the genotype [32]. Additional avian research in house sparrows suggests a role for changes in epigenetic potential (*i.e.,* the number of CpG motifs in the genome) during range expansions, while DNA methylation variation itself was inconsequential [33].

Hybrid zones are suitable natural models to decompose heritable genetic and epigenetic variation of diverged populations. In the central part of the hybrid zone, environmental variation is limited while genetic variation is maximized in mosaic hybrid genomes that are characterized by blocks of alternating ancestry. These properties have been successfully exploited to map the genetic basis of phenotypic variation and advance our understanding of the processes governing population divergence [34–36]. For studies of epigenetic variation, hybrid zones confer similar advantages [37]. We here make use of this fact and quantified both genetic and methylation variation in a well-studied avian hybrid zone.

All-black carrion crows (*Corvus (corone) corone*) and grey-coated hooded crows (*C. (c.) cornix)* hybridize in a narrow contact zone in central Europe which is governed by assortative mating and social dynamics related to plumage pigmentation patterns [38]. Genome scans have found minimal genetic divergence across most of the genome with the notable exception of a ∼2-Mb region on chromosome 18, hereafter referred to as the *focal region.* This region is subject to divergent selection, has accumulated fixed differences between parental taxa and is mainly responsible for the striking phenotypic variation [36,39,40]. Recombination is strongly reduced maintaining linkage disequilibrium of ancestral genetic variation [36,41]. The genetic heterogeneity of the system where a largely homogenous genome-wide background is opposed to a diverged *focal region* known to be relevant for speciation provides a unique opportunity to gain insight into the determinants of DNA methylation variation within and between taxa.

## Materials and methods

### Overview of study design and multiple-experimental control

We pursued two sampling strategies. We raised wild-caught nestlings from pure parental populations in Germany and Sweden to adulthood in a common garden experiment (hereafter ComGar., **Fig 1A** and **Fig 1B**) and sampled additional nestlings across a hybrid zone transect in Central Europe (hereafter HybZon.; **Fig 1C**). The ComGar. was primarily designed to establish a baseline of physiological determinants of 5mC methylation (tissue, age, sex) and test for taxon differences under controlled conditions; whereas the HybZon. experiment is suited to study the effect of genetic ancestry and environmental variation under natural conditions in the wild. In conjunction with whole-genome-resequencing data, both approaches further address the effects of genome properties (chromosomal and genomic features) and genetic variation within and between taxa (D_XY_, F_ST_, haplotype diversity, Tajima’s D, Fu & Li’s D*). To quantify the intensity of 5mC methylation per CpG site, we generated reduced representation bisulfite sequencing data which allows for comparisons of orthologous sites with high read coverage between individuals (ComGar.RRBS mean and standard deviation CpG coverage: 31.2 ± 3.87x; HybZon.RRBS: 45.2 ± 4.88x; **Fig S1**). For the ComGar. we additionally generated whole-genome bisulfite sequencing data allowing for technical validation of results across the entire genome and identification of a pervasive number of regions associated with tissue-specificity (ComGar.WGBS: 15.9 ± 1.22x) (see **<u>Table S1</u>** for an overview of the dataset). After filtering, we retained a total of 699,363 (ComGar.RRBS), 820,661 (HybZon.RRBS) and 4,190,434 (ComGar.WGBS) high-quality CpG sites.

**Fig 1.**
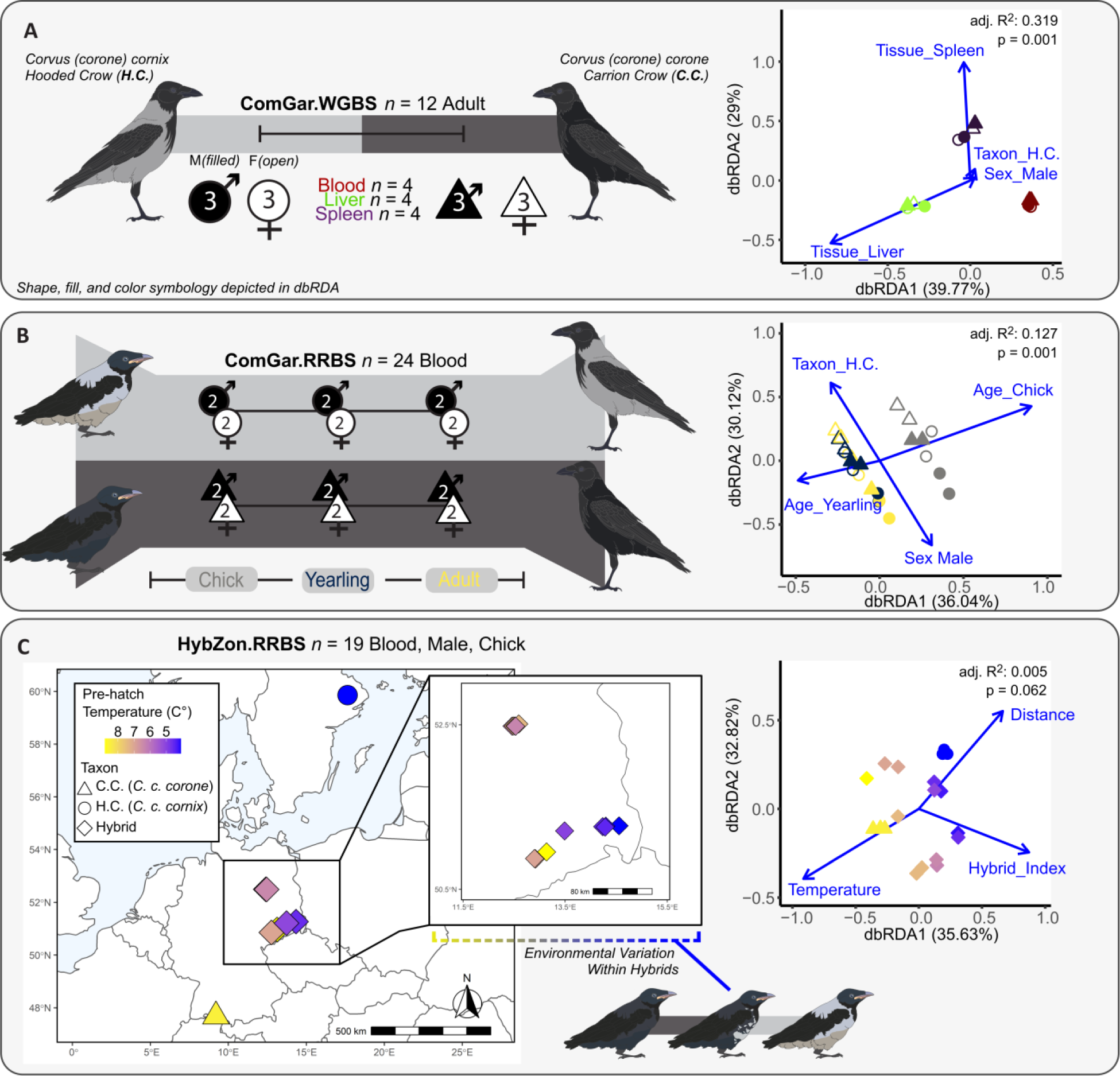
Experimental design to quantify the factors associated with 5mC methylome variation in crows. Left panels show sampling schemes for common garden (ComGar., A & B) and the hybrid zone transect (HybZon., C). (**A**) WGBS and (**B**) RRBS datasets to isolate the physiological effects of tissue, age, sex, and taxon. Number of post-filtering CpG sites (mean in millions): ComGar.RRBS: 0.699; ComGar.WGBS: 4.19. (**C**) RRBS data from transect-collected blood of male nestlings across the European hybrid zone (HybZon.RRBS: 0.821M CpGs) provides methylomic variation across environmental fluctuations (local pre-hatch winter temperatures) and genetic ancestry while controlling for tissue, sex and age. (**A - C**) Right panels show distance-based redundancy analyses (dbRDA) summarizing genome-wide DNA methylation variation within each experiment.

We used two approaches to assess the contributions of candidate variables (tissue, age, sex, taxon, environment) to methylation patterning: multivariate statistics examining global correlates of methylation variation and differential methylation analysis allowing for base-pair resolution of single differentially methylated positions. The multi-layered study design allowed us minimize false-positive inference of DMPs. For example, the ComGar.RRBS experiment contained chicks, yearlings and adults allowing identification of age-related CpG sites. We require that these age-related CpGs show no conflicting evidence from any of the other experiments (DMP for different variable, *e.g.* sex in the ComGar.WGBS) and, in addition, have low variance in the experiments that only considered a single age (*e.g.* ComGar.WGBS: all adults, HybZon.RRBS: all chicks). The latter condition excludes conflation with additional, non-measured variables. Last, we required a minimum effect size between groups (minimum 25% mean proportion difference). While allowing a margin for false negatives, this approach reduces spurious effects, which are rarely controlled for in studies of methylation variation [42]. Despite reduced power, it allows comparing the relative contributions of variables to methylation variation under the assumption of a similar effect size distribution.

### Sampling – Common Garden

For an illustrative overview of the sampling design see **Fig 1**. We designed a common garden experiment (ComGar.) to quantify DNA methylation variation corresponding to physiological factors (tissue, age, sex) and taxon differences. We first collected blood from the brachial vein of unrelated nestlings with a mean age of 22.1 days (range 14 – 26) from purebred carrion (*n* = 4) and hooded crows (*n* = 4). Carrion crows were collected in May 2014 from different nests in Konstanz in Southern Germany (47°45N′, 9°10′E) as part of a larger research program. Hooded crows were sampled around Uppsala, Sweden (59°52’N, 17°38’E) in the same month and year. These 8 individuals were transferred to Sweden by airplane and hand-raised indoors at Tovetorp field station, Sweden (58°56′N, 17°8′E). At the time when birds could feed independently, they were released into a roofed outdoor enclosure (6.5 × 4.8 × 3.5 m) and thereafter housed in single-sex groups of the same species under common garden conditions. For details of animal husbandry see [43]. Blood sampling was repeated for these 8 individuals during the non-reproductive season at an age of 18 months (551 – 565 days) and during the reproductive season at an age of 30 months when sexual maturity is expected to have been reached in all individuals (916 – 930 days) [44]. For this common garden experiment, sample sizes of both carrion and hooded crows raised in captivity were matched by sex (2 males and 2 females of each taxon) which was determined molecularly [45]. In addition to these 24 blood samples (8 individuals at three time points), we collected liver and spleen tissue from 4 of these individuals at sexual maturity, representing a male and female of each taxon (hooded and carrion crow) (**<u>Table S1</u>**). The 24 blood samples were subjected to RRBS (reduced representation bisulfite sequencing) to quantify sex, age, and taxon differences, while the 4 individuals with three tissue types available (blood, liver, spleen) were subjected to WGBS to quantify sex, tissue, and taxon differences (see below).

### Sampling – Hybrid Zone

We designed the hybrid zone experiment (HybZon.) to isolate genetic and environmental effects on DNA methylation variation while maintaining tissue, age, and sex constant. We sampled 3 purebred male chicks of each taxon (distinct from ComGar.) in addition to 16 hybrid chicks along a transect across the German hybrid zone during field trips in May 2008, 2013 and 2014. Hybrid individuals were characterized according to genotype information of 1,111 SNP markers spread across the genome that include two of the major loci coding for plumage colour variation, as outlined in Knief et al. [36] including a genetic factor on chromosome 18 (chr18) and the gene NDP on chromosome 1 with the R package introgress v1.2.3 [46]. Samples were chosen to represent the main diplotypes on chr18 (DD, DL, LL) and to encompass the full variation of genome-wide admixture (range of hybrid index: 0.0 – 1.0). With the inclusion of the purebred samples, hybrid indices thus both covered the full range from 0 (*C. c. corone*) to 1 (*C. c. cornix*) as represented by blood samples from 22 male individuals with an age range of 7 – 25 days (**<u>Table</u> <u>S1</u>**). In addition to genetic effects, this setup allows us to examine the influence of geographic distance and environmental variation. For the latter, we chose the mean temperature of the three months preceding hatch date, determined from Meteostat from the nearest local weather station (meteostat.net) (**<u>Table S1</u>**). This proxy for environmental effects thus integrates the maternal environment, as well as the environment experienced during the egg stage (incubation on average 18-19 days [44]).

### Sequence Data Generation - Methylome

We assessed genome-wide 5mC DNA methylation with a combination of whole-genome bisulfite sequencing (WGBS) and reduced-representation bisulfite sequencing (RRBS), involving four sequencing efforts. Genomic DNA isolated from whole blood, liver, or spleen using QuickExtract kits (Epicentre, Illumina) provided input for both WGBS library preparation (150 ng; *n* = 8 representing all tissues for each sex at parity) and RRBS library preparation (300 ng; *n* = 24 representing blood for three age classes of each sex and taxon at parity). Common garden experiment WGBS libraries were created following the TruSeq DNA Methylation kit (Illumina Inc., EGMK91324) according to the manufacturers’ protocol. RRBS libraries from the common garden experiment were created by adapter-ligating and end-repairing *MspI* digested fragments following the NEBNextUltra protocol (New England BioLabs). Bisulfite conversion was performed with the EZ DNA Methylation Gold Kit (Zymo) and bisulfite-converted DNA was then amplified with the NEBNext Universal primers and NEBNext index primers using 12 PCR-cycles. Libraries were cleaned twice using AMPureXP beads (55 ul beads to 50 ul sample). A 0.5% spike of non-methylated lambda phage DNA was included in WGBS and RRBS libraries to confirm bisulfite-conversion efficiency. Both WGBS and RRBS libraries were evaluated using a TapeStation with the HS D1000 kit. Adapter-ligated fragments were quantified with qPCR using a library quantification kit for Illumina (KAPA Biosystems/Roche) on a CFX384Touch instrument (BioRad) prior to cluster generation and sequencing. The WGBS libraries were then sequenced paired-end on a HiSeqX with 150-Bp read length using v2.5 sequencing chemistry, including a 5% PhiX spike-in. RRBS libraries were sequenced on a NovaSeq 6000 using an SP Flowcell, single-end 100-Bp read lengths using v1 sequencing chemistry, including a 25% PhiX spike-in. Sequencing was performed by the SNP&SEQ Technology Platform in Uppsala which is part of the National Genomics Infrastructure (NGI) Sweden and Science for Life Laboratory. In addition to this sequencing effort, we generated 22 RRBS libraries from the HybZon. experiment (19 surviving filtering, see below) and 4 supplemental WGBS biological samples (an additional two carrion crow tissues for each sex and supplemental sequencing for one individual from the previous WGBS effort) for the common garden experiment at a separate facility (Novogene, Co. Ltd.), bringing total sex, tissue, and taxon sampling to parity. RRBS *MspI*-fragmented libraries were created similarly to the common garden experiment detailed above, except the final libraries were sequenced paired-end on a NovaSeq 6000 with 150-Bp read lengths. The additional 4 WGBS libraries were generated at Novogene Co. Ltd. (Beijing, China), using the Accel-NGS® Methyl-Seq DNA library kit (Swift Biosciences) following the manufacturer’s instructions. These additional WGBS libraries were sequenced on a NovaSeq 6000 paired-end with 150-Bp read length. For treatment of batch effects see below.

### Genome Reference and Genomic Feature Annotation

All analyses were performed on the most recent chromosome-level European crow reference genome available at this time (National Centre for Biotechnology Information accession number: ASM73873v5). Analyses were carried out on all named chromosomes (*i.e.,* excluding unplaced scaffolds). Genomic features were extracted as follows. We annotated the reference genome into non-overlapping tracks of promoter regions, repeats, coding sequence (CDS), intronic, and intergenic sequence (**Fig S2**). Promoter regions were identified as CpG islands located within 2-Kb of a gene’s start coordinates. While this strategy will likely fail to identify many true promoters, particularly those with promoters not directly within the proximity of the transcription start site (TSS) [47], it will provide a conservative estimate of CpG islands with direct relevance to local transcripts. We created a *de novo* CpG island track using makeCGI v1.3.4 [48], requiring a length > 200-Bp, a GC content > 50%, and an observed/expected CG ratio > 60%, providing 30,459 islands. Only gene elements from the RefSeq .*gff* annotation file that intersected our CpG island track with the strand-aware 2-Kb region upstream of the gene start were retained, using bedtools v2.29.2 [49]. This resulted in 11,472 CpG-promoter islands, from a total of 17,944 genes. We identified repetitive regions with RepeatMasker v4.1.1 [50] using the repeat library from chicken and excluding simple repeats (‘-nolow’). Genic CDS and introns were extracted directly from the RefSeq annotation. A non-overlapping annotation feature track was created with bedtools and R v4.1.1 [51] by prioritizing promoter regions, repeats, CDS, introns, and intergenic annotations, in that order (**Fig S2**). To reduce erroneous RRBS alignments, we performed an *in silico* digest of the reference genome with *MspI* using SimRAD v0.96 [52], requiring a fragment size between 40 – 350-Bp, providing 299,587 potential fragments with roughly 1.6M CG motifs, compared to the roughly 9.8M CG motifs in the entire reference.

### Quantification of Genetic Variation

Genetic polymorphisms involving C-T and A-G transitions are problematic for bisulfite sequencing experiments because C-T and A-G mismatches between sequence and reference are used to identify DNA methylation [53,54]. We therefore exploited an existing population resequencing dataset from 28 male hooded and carrion crows sourced from the same allopatric populations (Uppsala, Sweden and Konstanz, Germany) analyzed in this study (*n =* 14 of each taxon) to identify transition SNPs. We also used this dataset to quantify genome-wide genetic variation (F_ST_, D_XY_, Tajima’s D, haplotype diversity, Fu & Li’s D*) to assess its relationship with methylation variation. In short, 130 paired-end Illumina libraries for the 28 samples were adapter-trimmed with BBTools v38.18 [55], aligned to the reference with BWA v0.7.17 [56], merged with samtools v1.7 [57], and deduplicated with GATK v4.1.9.0 [58]. After read trimming and filtering, alignment to the hooded crow reference genome (accession ASM73873v5) resulted in a genome-wide mean coverage of 15.1x (range 8.03 – 30.9x), calculated in 100-Kb windows with mosdepth v0.3.1 [59] (**<u>Table S2</u>**).

An all-sites variant file was called with bcftools v1.16 [60]. For filtering variant sites, we retained only biallelic sites passing hard filter thresholds (‘QUAL > 20, DP < 2*Average DP, or DP > 28x’), which had scored genotypes in at least 90% of individuals (*n =* 25/28 with *FMT/DP >= 3*). Following hard site-level and genotype-level filters (**Fig S3** and **Fig S4**), we retained 9.44m variant sites and 1,010m invariant sites. Coordinates for transitions (C-T and A-G polymorphisms) were subset using bash scripts for methylation site masking. Population genetic metrics for hooded and carrion crows (F_ST_ and D_XY_ between taxa and Tajima’s D, haplotype diversity, and Fu & Li’s D* among all individuals) were calculated in windows of 5000 sites (‘--windType sites --windSize 5000’), requiring a minimum of 1000 shared sites across samples (‘--minSites 1000’) using the genomics_general repository [61] for F_ST_ and D_XY_, and the *Theta_D_H.Est* repository for the remaining metrics [62]. Principal components analysis (PCA) on LD-pruned genome-wide SNPs was performed with SNPRelate v1.28.0 [63], and again exclusively on the SNPs falling within the *focal region* on chromosome 18. Genetic data were visualized with karyoploteR v1.20.0 and ggplot2 v3.3.6 [64,65].

### Quantification of 5mC Methylation

All bisulfite sequence datasets were first trimmed with trim_galore v0.6.7 [66], ensuring the artificial filled-in positions were trimmed in the RRBS datasets (‘--rrbs’), and requiring average read Phred scores of at least 20 (‘--quality 20’). Bisulfite conversion was successful, measured using a non-methylated lambda phage DNA spike into each library (minimum, mean, maximum conversion rate: 98.90, 99.27, 99.74%, **<u>Table S3</u>**). Reads were aligned to the European crow reference genome using Bismark v0.23.0 with default settings [53]. We extracted CpG methylation calls using Bismark, iteratively repeating this process after ignoring read positions exhibiting systematic methylation biases, evidenced through methylation bias plots (**Fig S5**). WGBS post-bisulfite adapter tagged (PBAT) libraries required additional trimming before alignment to remove artificial bases inherent to the protocol (‘--clip_r1 9 --clip_r2 9 --three_prime_clip_R2 1’). We then retained methylation calls on *MspI* fragments (for RRBS data only), removed any sites overlapping a C-T or A-G SNP (identified from the genetic resequencing experiment, see above) using bedtools v2.29.2 [49] or located on a scaffold (as we only analyzed sites on assigned chromosomes), and removed any sites covered by less than 10 reads in each individual for the RRBS experiments, and less than 5 reads for the WGBS experiment. We chose a lower threshold within the ComGar.WGBS experiment to maximize retained sites. RRBS reads were ensured to overlap an *in silico* digested *MspI* fragment (see above) using bedtools [49]. Initial count inspections for RRBS data indicated a substantial increase in retained reads when overlapping *MspI* fragments at the BAM file level (pre-methylation extraction), as opposed to *post-hoc* removal of individual sites after extraction, so extraction on RRBS dataset was repeated after filtering BAM files for *MspI* fragments. Methylation levels (%) were consistent across read positions after trimming, indicating no systematic positional read biases, although high coverage spikes in the RRBS datasets lead us to set upper coverage limits on our datasets, for which we removed a conservative degree to alleviate any concerns with repetitive fragments that would be difficult to map with A-T rich bisulfite sequencing reads (remove the top 5% coverage outliers across all experiments). Finally, CpGs were removed from the experiment if they had missing data from any individual (ComGar.WGBS: *n =* 12/12), more than 3 individuals (ComGar.RRBS: *n =* 21/24), or 1 parental and 2 hybrids (HybZon.RRBS: *n* = 16/19) to ensure that sufficient replicates for each biological treatment existed for each site.

No batch effects associated with the two sequencing centers were apparent in the ComGar.WGBS experiment (**Fig S6**), identifying very repeatable measurements across sequencing centers (Spearman’s rho: 0.998 for all shared CpGs above 10x coverage). Initial ordinations of the HybZon.RRBS experiment indicated three hybrid individuals (Y13, Y31, Y40) as outliers, seemingly due to either systematic hypomethylation and/or age in one sample (**Fig S7**). To be conservative we excluded all three individuals, as no known covariates were immediately responsible for driving the variation in all three individuals and they did not contribute to novel axes of *a priori* physiological variation (*i.e.,* the full range of hybrid indices were still retained). We replicated DNA extraction, library preparation, and sequencing for four biological samples for both RRBS and WGBS protocols, indicating higher repeatability than expected by chance (**<u>Fig</u> <u>S8</u>**). While correlations were lower than between technical replicate WGBS libraries (see above), these differences are likely explained by inherent differences between these protocols, sequencing coverage, and blood as a source tissue which may contain different cellular make-up in each extraction. Hatchlings were sampled from different nests to ensure unrelated individuals were sampled, and extra pair paternity in crows is low [67].

### Determinants of Methylome Variation

The main goal of this study was to examine the variables associated with variation in methylation. Variables under investigation can be grouped into genome properties, genetic variation, physiological determinants, and environmental contribution. Depending on the variables under investigation, different approaches were used (multivariate statistics, beta-binomial regression, supervised machine learning). We examined i) the global determinants of physiological and environmental methylation variation with constrained ordinations; ii) base-pair resolution divergence associated with physiological, environmental, and genetic effects (*i.e.,* taxon, hybrid index) with beta-binomial regressions; and iii) genome property and genetic variation covariation with DNA methylation using supervised machine learning and bootstrapped correlations. Methylation data were treated according to the method requirement as a methylation proportion (ranging from 0.0 – 100% for distance-based redundancy analyses and machine learning regressions), or directly as methylated and non-methylated read counts (for DMP detection with beta-binomial regressions). We ensured analytical insensitivity by repeating analyses, where applicable, using multiple methylation inputs (*e.g.,* we analyzed global variation with unconstrained ordinations using both Euclidean and alternative distance measures; PCA and NMDS; analyzed the machine learning problem as a classification problem instead of a regression using binary and trinary discrete methylation response variable categories).

### Physiological and Environmental Determinants

To assess the effects of tissue, age, sex, taxon, and environment on DNA methylation patterns, we used two approaches: multivariate statistics examining global correlations of methylation variation (distance-based redundancy analyses; dbRDA), and differentially methylated position (DMP) analyses allowing for ultra-fine resolution of single differentially methylated CpGs. First, for each experiment independently (ComGar.WGBS, ComGar.RRBS, HybZon.RRBS), we examined global DNA methylation variation with a dbRDA constrained ordination. Methylation input for the ordination was a matrix composed of DNA methylation proportions with columns corresponding to each individual CpG site and rows corresponding to individual samples. Optimal distance measure for dbRDA (and later NMDS) for each experiment was identified using Spearman’s rank correlations within vegan v2.5-7 using the rankindex function (Gower’s distance: ComGar.WGBS; Euclidean distance: HybZon.RRBS, ComGar.RRBS). A pre-processing data recipe was created with tidyverse v1.3.1 and tidymodels v0.1.4 [68,69], ensuring variables were in proper classes and that collinearity between variables was below a Spearman’s rank correlation of 0.8. Scaled dbRDAs were then implemented with vegan, and the contributory effects of each variable were assessed with an ANOVA by term with 10,000 permutations (**<u>Table S4</u>**). Adjusted R^2^ and overall model significance as assessed with an ANOVA are reported. To minimize statistical biases inherent in any single method [70], we also performed an unconstrained ordination (NMDS) including an analysis of similarity (ANOSIM) on DNA methylation variation and a typical eigenvector based analysis (PCA; **Fig S9** and **<u>Table S4</u>**), and observed the same patterns.

We then determined the physiological and environmental determinants of DNA methylation at CpG-resolution using DMP detection directly from methylated and non-methylated read counts. Explanatory variable effect estimates (test-statistics) and FDR-corrected *p*-values were obtained for each variable and for each experiment independently using a beta-binomial regression with an arcsine link implemented in the R package DSS v2.42.0 [71]. Explanatory variables for ComGar.WGBS data were tissue (Blood, Liver, Spleen), sex (Male, Female), and taxon (C.C., H.C.); for ComGar.RRBS were age (Chick, Yearling, Adult), sex (Male, Female), and taxon (C.C., H.C.); for HybZon.RRBS were genetic ancestry hybrid index, geographic distance, and mean local temperature for the three months preceding hatch date (*i.e.,* a proxy for environmental effects), determined from Meteostat from the nearest local weather station (meteostat.net) (**<u>Table S1</u>**). An additional variable of sampling year was also investigated, but the patterns appeared well summarized with the existing variables (**Fig S10**). We then assessed taxonomic differences in DNA methylation within the focal region compared to the autosomal background using bootstrapped sampling with replacement within R [51]. For each DNA methylation experiment and each genomic annotation feature, we sampled a number of autosomal CpGs equal to the number of CpGs within the focal region for that subset and calculated the mean taxon effect estimate, repeating this process 1,000 times. We considered the comparisons to be non-overlapping if the 95% quantile distributions did not overlap.

### Multi-experimentally Verified DMPs

Within each experiment independently, individual CpG sites were then assigned a corresponding physiological or environmental classification (*e.g.,* tissue, age, sex, taxon, temperature, distance), or were marked as conserved (less than 10% variation within entire experiment), or indeterminate (no significant variation corresponding to any of the covariates). Classifications were based on an FDR detection threshold below 10%, non-significance for other covariates (FDR > 10%), and a conservative minimum divergence between group methylation means of variables of interest of 25% (*e.g.,* absolute value of mean(Male) – mean (Female) > 25%). Concordance between FDR-corrected *p*-values and methylation divergence for each effect were checked with volcano plots (**Fig S11**). No absolute divergence thresholds were used for continuous covariates (hybrid index, temperature, geographic distance) as these were not factor variables and did not contain groups. Conserved sites represented a small amount of CpGs within ComGar.WGBS (10.3%, *n =* 432,403), and higher relative amounts in the promoter-rich reduced-representation experiments HybZon.RRBS (49.7%, *n =* 407,481) and ComGar.RRBS (48.5%, *n =* 339,410).

Classified CpGs were then merged across experiments at base-pair resolution and subjected to multi-experimental control. A classification was considered verified for a site if i) it was inferred in at least one experiment, ii) there was no conflicting classification among all three experiments, and iii) at least one experiment serving as a negative control classified the site as conserved (***cf.* Fig 2**). For instance, for a verifiable age DMP classification the site would have to be classified as age-specific in the ComGar.RRBS and be conserved in either ComGar.WGBS or HybZon.RRBS (or both), where age is controlled for. As we had no independent experiment with a negative taxon control, we required significance for hybrid index within the HybZon.RRBS experiment, as well as one corroborating taxon classification from the other experiments. While this approach relies on CpG overlap between experiments, which is of course limited by coverage and technical conditions (*i.e.,* we expect a high number of sites present in WGBS to be absent in RRBS), we preferred this strategy as it will dramatically reduce false positives by introducing multi-experimental reproducibility within a single study. Furthermore, as our primary interest is on DNA methylation divergence between taxa, integrating two metrics of taxonomic divergence (*i.e.,* parental comparisons in two experiments and hybrid index in another) will identify sites verifiably corresponding to taxonomic divergence. While financial constraints may cause our study to miss small-effect DNA methylation loci due to sample size limitations, our sample size within each experiment matches those in other WGBS [72–76] and RRBS [77–81] studies. Our multi-experimentally validated approach offers a novel framework to assess both inter-experimental batch effects and repeatability in a conservative framework, and our preferred approach in contrast to an approach which maximizes DMP discovery at the cost of false positives.

**Fig 2.**
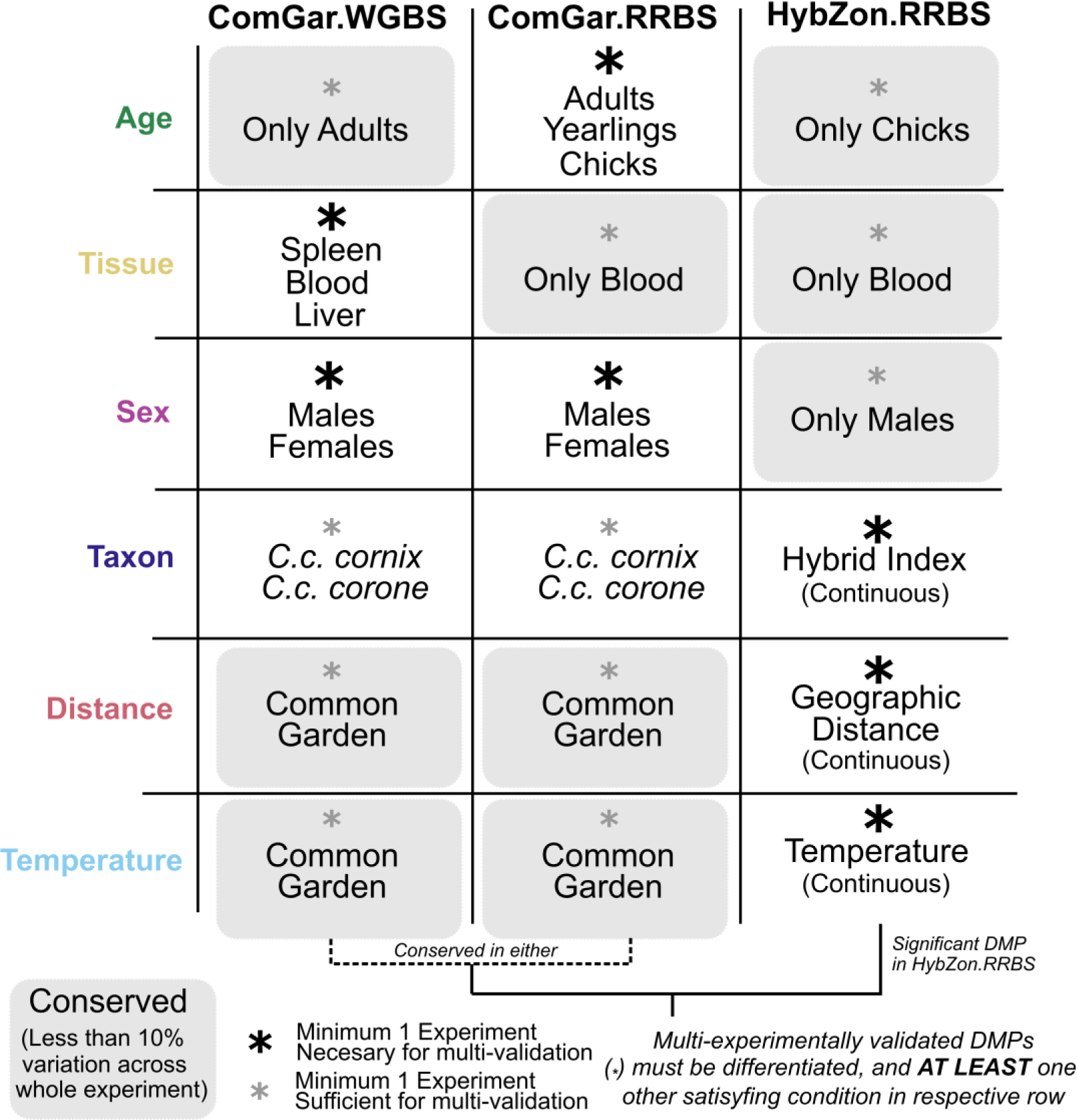
Multi-experimental validation to remove spurious DMPs. Each sequencing effort includes unique biological variation that can be exploited to control for false-positive inference (*i.e.,* multi-experimental support). Differentially methylated positions (DMP) are considered valid only if the site shows 1) statistical evidence for differential methylation in at least one of the corresponding asterisked fields in its row (*necessary condition, black asterisk*), 2) no conflicting evidence from any different variable and 3) support from *at least* one of the other classifications in that row (*sufficient condition, grey asterisk*). Sufficiency requires support for the variable of interest in another experiment (*e.g.* taxon) or low variation in an experiment where this variable has only single-state (*e.g.* age) exhibiting less than a 10% variation in methylation (*conserved, grey shading*).

We assessed sensitivity in our classification strategy outlined above by repeating the process except classifying sites based solely on the common garden experiments (*i.e.,* not requiring significance for any effects in the HybZon.RRBS). We then overlapped the HybZon.RRBS CpGs with these single-experimentally classified sites and examined the distributions of methylation variation within sites classified as *Conserved, Tissue*, or *Taxon* from the ComGar. We examined methylation variation within each hybrid group: unadmixed hooded and carrion crows, hybrids which have hybrid indices (calculated using the chromosome 18 and NDP factors) completely hooded or carrion (either 0.0 or 1.0), and hybrids intermediate to either. Permutation tests between methylation levels within the focal region and compared to the autosomal background corroborated the results found within **Fig 3D**, showing hypomethylation within carrion crows and their closely-related hybrids within the island on chromosome 18, and hooded crows and their associated hybrids hypermethylated compared to the autosomal background (**Fig S12**). Similarly, hybrids with intermediate genetic ancestry coefficients exhibited intermediate methylation levels which did not differ from the autosomal background. No differences were observed between the focal region and the background within *Conserved* CpGs, which closely mirrored the tight distributions seen for *Tissue* windows, although we did observe constitutive hypomethylation within the focal region for tissue windows within the HybZon.RRBS (**Fig S12**).

**Fig 3.**
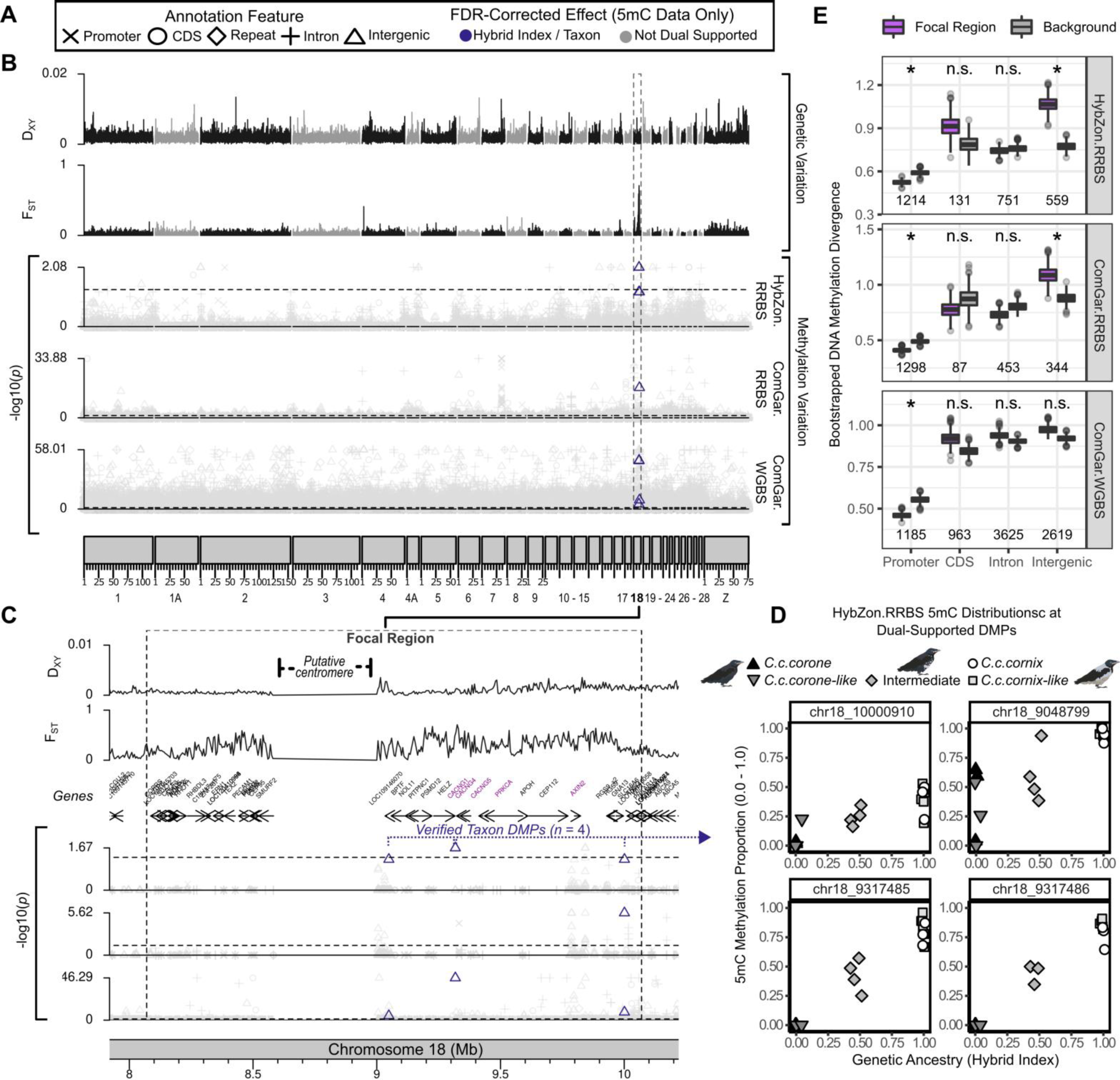
Methylomic variation along the crow genome. (**A**) Legend used throughout describing genomic annotation feature (symbol) and explanatory variable (colour) for each DMP. (**B**) *Genetic variation:* genetic divergence (D_XY_ (minimum, mean, maximum): 0, 0.0015, 0.013) and genetic differentiation (F_ST_: 0-0.01-0.71) across all *n* = 28 allopatric hooded and carrion crows. The region of elevated differentiation on chromosome 18 (focal region) is highlighted (99% of sites with F_ST_ > 0.3). *Methylation variation*: FDR-corrected base-pair level *p*-values with coloured points indicating taxon-specific differentially methylated positions for multi-experimentally verified DMPs. Only the lowest FDR-corrected effect was plotted for each CpG in each experiment. To aid in visualization a ceiling is imposed on minimum and maximum values within each experiment based on quantiles of the limits of visible data (*see methods*). (**C**) The *focal region* of high genetic differentiation (F_ST_ = 0.298 within vs. 0.010 outside) shows the only four multi-experimentally verified DMPs related to taxon divergence. Candidate genes underlying plumage polymorphism from Knief et al., 2019 are highlighted in purple. Otherwise, tracks as in panel B. (**D**) Methylation levels (y-axis) closely follow genetic ancestry (x-axis) at the four taxonomically-related sites following the general pattern of hypomethylation of carrion crow ancestry in this region (**Fig S12**). (**E**) Methylation divergence for all CpGs within the focal region compared to the autosomal background. Default parameter boxplots correspond to first and third quantiles; significance indicated if either side of the 95% distribution tails do not overlap. Total CpGs used for empirical counts within focal region shown below box plots.

### Genome Properties and *cis*-genetic Variation

Following the observation that all (*n =* 4, **<u>Table S5</u>**) multi-experimentally verified taxa DMPs were within a region of elevated genetic variation on chromosome 18, we sought to quantify the relationship between taxonomic DNA methylation divergence and its chromosomal substrate at genome-scale, namely using genome properties and population genetic variation. We used the taxon-related (*e.g.,* taxon for ComGar. or hybrid index for HybZon.) test-statistics from each experiment’s DMP analyses as a response variable input, and intersected these with the population genetic variation (F_ST_, D_XY_, Tajima’s D, haplotype diversity, Fu & Li’s D*), and genome property information (genomic annotation, GC-content, relative position of the window along a chromosome, and chromosome length. Population genetic variation was assessed genome-wide in windows of 5-Kb, so we calculated mean DNA methylation divergence (taxa-related test-statistics from DMP analyses) within each window. For sensitivity, we also repeated the entire following analysis with maximum divergence values within 5-Kb windows instead of the mean and arrived at similar conclusions, except that the importance for *promoter* regions was relinquished into GC-content, highlighting the interconnected nature of these covariates (**Fig S13**). We imputed population genetic values from the nearest 5-Kb windows where values did not exist (mean missing 5-Kb windows across experiments: *n* = 240, 0.78%), seemingly more common in areas of high GC-content (within missing windows = 68.0%, outside = 57.3%; **<u>Table S6</u>**). We then assessed the raw distributions of our DNA methylation divergence response variable within each experiment and deemed a log transformation appropriate (**Fig S14**). Collinearity between explanatory variables was low across all three experiments (minimum, mean, maximum: −0.27, 0.056, 0.62; **<u>Table S7</u>**).

We assessed the relationship between DNA methylation divergence and its chromosomal substrate using supervised machine learning regression (*i.e.,* random forests with ranger v0.14.1 [82] and boosted regression trees with XGBoost v1.7.1.1 [83]). Pre-processing and modeling used a tidyverse v1.3.1 and tidymodels v0.1.4 [68,69] framework implemented within R v4.1.1 [51], where numerical covariates were normalized and collinearity between variables was checked (Spearman’s rank correlation < 0.8). Datasets were then stratified into 75% training and 25% testing partitions to be analyzed with 5*-*fold cross validation. Model parameters (‘trees, min_n, mtry’; and ‘learn rate’ for XGBoost) were tuned to maximize R^2^ with finetune v1.0.1 and caret v6.0-93 [69,84] using a random grid search. Workflows were finalized and implemented with tidymodels using the tuned parameters, allowing for final extraction of covariate permutation importance using vip v0.3.2 [85] and model fit (R^2^ and root mean squared error, RMSE) (**<u>Table</u> <u>S8</u>** and **<u>Table S9</u>**). Permutation importance measures explanatory variable importance by shuffling each covariate’s values and measuring the impact on model fit, and is a better gauge of contributory effects than methods relying on impurity [86]. Permutation importance is therefore a useful measure to determine the importance of each covariate, but only insofar as the final models has sufficient overall explanatory power. To generate confidence intervals on covariate importance and model explanatory power, we replicated the entire process three times for all three DNA methylation experiments using different random seeds. We then scaled permutation importance within each iteration to a value of 1.0 to allow for comparisons across the two regression engines. Furthermore, to see if the problem could be approached more parsimoniously with supervised classification techniques, we transformed our continuous DNA methylation divergence response variable into a discrete binary (and trinary) response and repeated the entire process again (**Supplementary Text, Fig S15**), this time assessing models with the Area Under the Receiver Operator Characteristic curve (ROC AUC) and covariate permutation importance, and observed the same patterns **(<u>Table S8</u>, <u>Table S9</u>,** and **Fig S16)**.

We complemented our supervised machine learning approach with a bootstrapped Spearman’s rank correlation analysis between taxonomic DNA methylation divergence and population genetic variation, with a focus on how patterns differ between the focal region on chromosome 18 and the autosomal background. The boundaries of the focal region were established by the start and end of windows on chromosome 18 with elevated genetic differentiation (F_ST_ > 0.3). At this F_ST_ threshold nearly all windows were localized on chromosome 18 (99.0%, *n =* 149), and the resulting region (starting at 8.07e6 and ending at 10.07e6, 2-Mb) corresponds to the length of the island identified previously, although on a different genome assembly [36,87]. For each DNA methylation experiment, we sampled (with replacement) autosomal windows equal to the number of focal windows and calculated Spearman’s rank correlation between DNA methylation divergence (absolute value of test-statistic estimates for taxon effects from DMP analyses) and population genetic measures inside and outside of the focal region. We considered the correlations to be significant if their 95% quantile distributions did not overlap zero.

## Results and discussion

### Ontogenetic program

First, we assessed the physiological determinants contributing to the organism’s ontogenetic program. Tissue-specificity dominated DNA methylation variation in the ComGar.WGBS experiment (dbRDA *p*-value for tissue effects; *p_(Tissue)_* < 0.001). Tissue explained a large proportion of overall methylation variation along with sex and taxon (overall dbRDA; *p* < 0.001, adj. R^2^ = 0.32), each having little effect by itself (*p_(Sex)_* = 0.11, *p_(Taxon)_* = 0.096) (**Fig 1A** and **<u>Table S4</u>**). Within the ComGar.RRBS experiment, age class was the primary driving force. Cumulatively with sex and taxon, age stage between chick and mature crows explained moderate variation in methylation patterns (adj. R^2^ = 0.13, all 3 variables *p* < 0.001), while there was no detectable difference between yearlings and adults (*p_(Yearling)_* = 0.97) (**Fig 1B**). Within the HybZon.RRBS experiment, continuous proxies for taxon (hybrid index), geographic distance, and environmental variation (pre-hatch ambient temperature) were nearly uncorrelated with methylation patterning (adj. R^2^ = 0.005), and only temperature was identified as significant (*p* = 0.008) (**Fig 1C**). Ambient temperatures during development are known to impact global methylation patterning in birds [88], so our results provide some support for temperature regulation of the DNA methylation program, although the effects are slight.

At CpG resolution, DMPs largely mimicked results from the global analyses, but also provided the basis to pinpoint the genomic regions associated with taxonomic divergence after controlling for other effects of the ontogenetic program (**Fig S17**). By far the most well-supported DMPs were related to tissue (*n =* 3,863) followed by sex (*n =* 6), age (*n =* 4), taxon (*n =* 4), pre-hatch ambient temperature (*n* = 3) and geographic distance (*n* = 1) (**<u>Table S10</u>**). Note that the low numbers of DMPs are a consequence of multi-experimental control minimizing false-positive inference at the cost of false negatives (**Fig 2**). Since multi-experimental control is predicated on CpG overlap across experiments, the total number of usable CpGs was substantially reduced to 2.11% (*n* = 105,559) sequenced across all 3 experiments and 12.02% (*n* = 601,516) in at least two experiments (**<u>Table S11</u>**). Within the latter category, 63,073 CpGswere classified as DMPs satisfying statistical significance from beta-binomial regressions and an effect size threshold of 25% difference in methylation (necessary condition **Fig 2**). Of these putative DMPs, 872 (1.38%) showed conflicting classifications across experiments and were accordingly dropped (*e.g.* a taxon effect in one experiment, and an age effect in another). Of the remaining 62,201 only 6.24% (*n* = 3,881) survived the sufficiency criteria of multi-experimental control (**Fig 2**, **Fig S18**, **<u>Table S10</u>** and **<u>Table S12</u>**).

In summary, the results on physiological determinants implicate overarching DNA methylation regulation primarily in cell fate and age-related processes, reinforcing the well-established role of DNA methylation in cell lineage differentiation [89,90] and ageing [91]. Ancestral genetic variation separating the young taxa had a substantially less prominent signature genome-wide (**<u>Fig</u> <u>S19</u>**).

### Methylomic and genomic interplay

The effectsof genetic ancestry on methylation patterns were restricted to the focal region of genetic differentiation primarily responsible for plumage polymorphism. All four robust taxon-related DMPs were restricted to this region and showed evidence for hypomethylation of carrion crow ancestry (**Fig 3C**, **Fig 3D, Fig S12, Fig S20, Fig S21, Fig S22, Fig S23, <u>Table S13</u>**). Encoding of methylation genotypes with k-means clustering at these loci identified strong methylation linkage disequilibrium between these sites (D = 0.185, D’ = 0.743) (**Fig S24)**, mirroring phenotypically-associated linkage blocks identified from genetic data in this region [36]. The fact that these taxonomically relevant DMPs all occur in proximity to the plumage polymorphism candidates AXIN2, PRKCA, and an array of CACNG genes [36] support the notion that phenotypically relevant genetic variation and DNA methylation may indeed covary as a result of divergent selection, as has been proposed in white-throated sparrows [92] [but see 26]. DNA methylation may thereby act as mediator translating signals of *cis-*genetic variation into differential physiological activity. Note, however, that the taxonomic DMP loci were identified using tissue (blood, spleen, liver) which is not histologically relevant to melanin production in crows [93], and would therefore reflect a tissue-unspecific, pleiotropic signal. Moreover, none of the DMPs overlap any genes or promoters directly, and more generally, elevated methylation divergence was restricted to intergenic space while it was decreased in promoters compared to the autosomal background (*p* < 0.05 in all experiments; **Fig 3E, <u>Table S14</u>**). Altogether, this renders functional importance of the 5mC DMPs in the focal region less likely.

Having established the effect of local genetic divergence on 5mC methylation, we next assessed whether taxonomic variation in DNA methylation could be predicted more broadly. Using supervised machine learning regressors, methylation divergence between taxa (**Fig 4A**) was in part predicted by genome properties (chromosome length, positioning along chromosome, functional annotation) and to a lesser extent by measures of population genetic variation (F_ST_, D_XY_, haplotype diversity, Tajima’s D, Fu & Li’s D*) (**Fig 4B** and **<u>Table S8</u>**). GC-content (Permutation Importance 95% CI intervals across all experiments: 19.4 – 24.0%) and promoter regions (16.9 – 27.6%) were the strongest predictors indicating decreased methylation divergence within GC-rich CpG islands near putative promoter regions (**Fig 4C, <u>Table S8</u>** and **<u>Table S9</u>**). This finding corroborates the prevalent conservation of evolutionary processes governing CpG islands in vertebrates [94,95]. Relative positioning along a chromosome (4.35 – 5.91%) and chromosome length (5.78 – 8.22%) were weaker predictors of taxon methylation divergence, yet lend support for methylome divergence proceeding more rapidly on micro-chromosomes or near chromosome ends where recombination is substantially higher [96]. Further supporting the relationship between genetic variation and DNA methylation, permutation importance was highest for a measure of genetic divergence, D_XY_ (8.63 – 14.7%), followed by haplotype diversity (9.00 – 13.4%), Tajima’s D (5.62 – 8.87%) and eventually genetic differentiation (F_ST_ 1.38 – 2.74%) (**Fig 4B** and **<u>Table S9</u>).** Our results indicate interplay between population genetic variation and DNA methylation and supports previous research identifying population-level correlates between genetic and DNA methylation variation [24,97], particularly in regions undergoing selection [98]. To compare the relative strength and obtain directionality of the interplay between DNA methylation and population genetic variation, we examined bootstrapped correlations within the focal region compared to the autosomal background. As expected, the relationship between DNA methylation and genetic variation was stronger within the focal region, particularly for D_XY_ (**Fig 4C** and **<u>Table S15</u>**).

**Fig 4.**
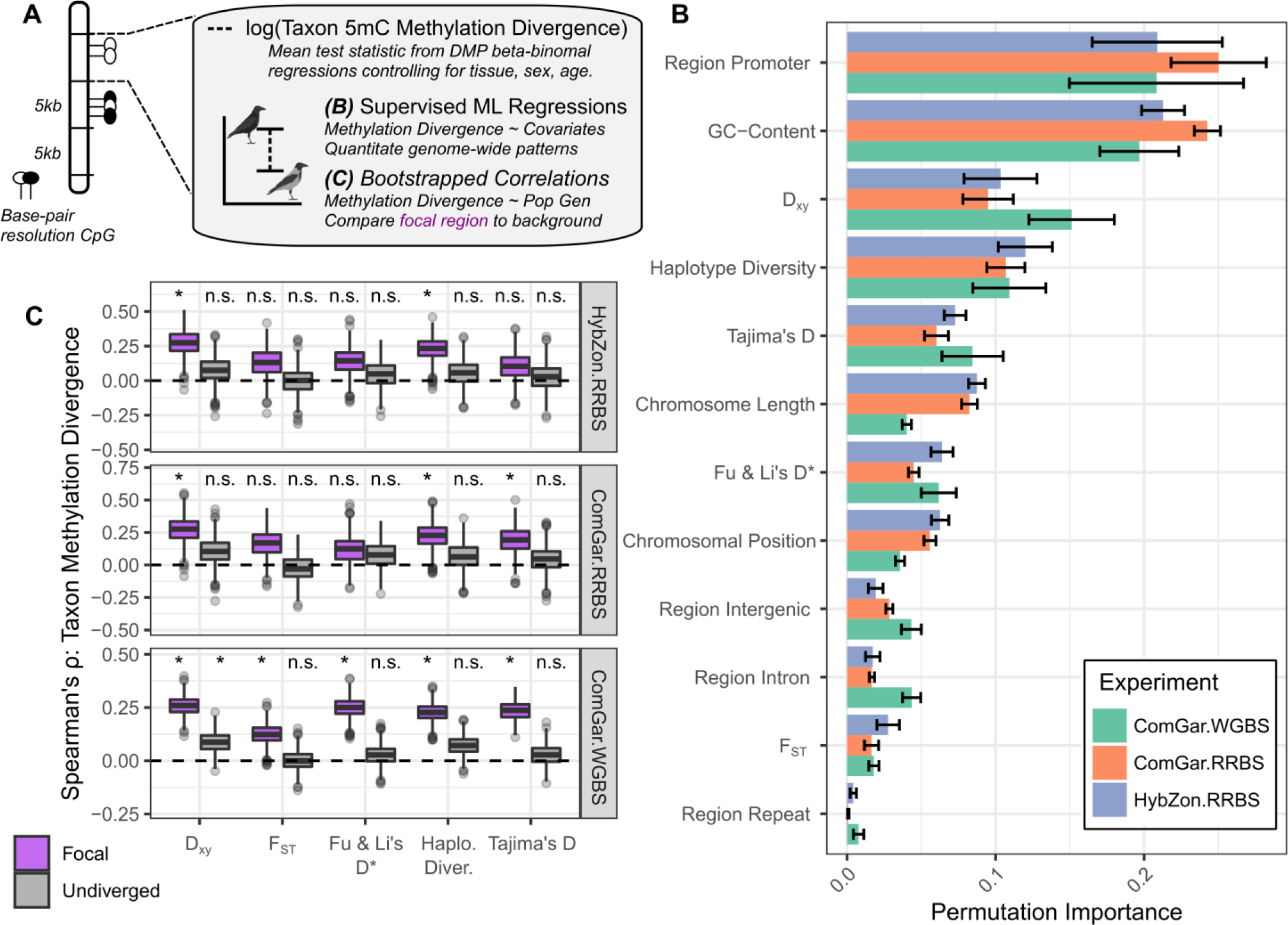
Predicting 5mC methylation divergence between hooded and carrion crows. (**A**) Cartoon depicting the two methods used to examine broad chromosomal and population genetic patterns associated with DNA methylation. (**B**) A supervised machine learning regression approach across the three methylation sequencing efforts determined the relative contributions of explanatory variables (y-axis) to taxonomic methylation divergence. Methylation divergence was summarized as the binned mean of DMP output test-statistic estimates for taxon-effects, controlling for other variables within each experiment (**Fig 2, Fig 3, Fig S21**). Permutation importance measures variable importance against a decrease in model fit and indicates predictive ability. Mean R^2^ across each experiment was 0.050, 0.097, 0.21. (**C**) Spearman’s rank correlation between DNA methylation divergence and population genetic metrics within and outside of the focal region, indicating elevated covariation within the focal region which is characterized by a higher degree of genetic differentiation. Distributions drawn from 1,000 bootstrap sampling events; default parameter boxplots correspond to first and third quantiles; significance indicated if either side of the 95% distribution tails do not overlap zero

In summary, genome properties and segregating genetic variation both contribute to taxonomic divergence of DNA methylation, most pronounced in the *focal region* undergoing divergent selection. These results are overall consistent with a general carry-over effect of *cis-*acting genetic variation on patterns of methylation variation, as observed within *Papio* baboons [99]. Hitchhiking 5mC variants in linkage disequilibrium with *cis-*acting genetic variation may or may not be co-opted functionally [6,100–102] (for an example of *trans-*acting genetic variation see [103]). Conversely, linkage disequilibrium could be reinforced by selection on DNA methylation, dragging along genetic variation. The known causal effects of DNA methylation on genetic variation are currently limited to increased deamination of methylated cytosines [104], with these effects most pronounced in TEs [105]. While the gain or loss of genomic CpG motifs via deamination from DNA methylation could have evolutionary implications [33,106], this conclusion is less parsimonious in our system where there is no evidence for divergence in TEs between taxa [107] and intergenic taxonomically relevant DNA methylation is restricted within the large 2-Mb *focal region* responsible for phenotypic divergence between taxon [38].

### Environmental Effects

Environmental inducibility of epigenetic variation has received much attention [108], but data under natural conditions is scarce and near-absent in animals [26,28]. The fact that environmental contrasts generally covary with genetic effects further complicates the quest [109]. Our experimental setup allowed us to separate these and other confounding variables. To isolate environmental effects on DNA methylation variation, we focused on the HybZon.RRBS experiment which excludes all physiological confounding variables *a priori* (all blood from male chicks). We then isolated any conserved sites which exhibit less than a 10% range in DNA methylation in the ComGar. experiments, but show significant effects for pre-hatch temperature in the HybZon.RRBS experiment. Incubation and early developmental ambient temperatures are known to effect DNA methylation patterns in numerous tissues in broad avian taxa ranging from chickens [110], duck [111], and zebra finch [88]. While ambient temperature has been implicated in DNA methylation patterns related to reproductive timing in great tits [27,112], our study supports the notion that genetic effects impart more to the epigenetic program than early rearing environment, as indicated by cross-fostering experiments [32]. While a standard experimental approach would have identified numerous environmentally-sensitive loci (*n =* 61), our multi-experimental approach revealed only 3 verifiable DMPs related to ambient pre-hatch temperatures (**Fig S17** and **<u>Table S5</u>**), indicating hypervariable loci or off-target effects may be responsible for numerous candidates in studies examining environmental patterns in wild populations. One of our multi-experimentally validated DMPs lies within the promoter of *LMF1*, a gene whose methylation status in humans is associated with offspring birthweight [113]. While it lacks any functional validation in avian systems, its implication in gene-environment associations during embryonic development [114] indicates the methylation status of *LMF1* as a candidate associated with early developmental environment. The remaining two DMPs lie in the vicinity of long non-coding RNAs. Using sampling year as an additional measure for environmental variation yielded no multi-experimentally validated DMPs and recovered the same patterns associated with temperature (**Fig S10**). While these results do not provide mechanistic insight, they do not preclude a possible role of DNA methylation in mediating environmental stimuli to regulatory gene activity irrespective of underlying genetic sequence [27,115].

Overall, this study illustrates the utility and limitations of methylomic approaches and advocates a multiple, hierarchical experimental framework for addressing complexities that underlie genetic-epigenetic-environmental interactions in natural populations. While the experimental design did not allow assessment of the extent of spontaneous, putatively heritable epigenetic mutations, it provides clear evidence that natural variation in DNA methylation is firmly associated with general genome features and physiological processes orchestrating cell fate and aging. The latter conforms to the original definition of epigenetics as ‘the interaction of genes and their products […] which bring the phenotype into being’ [5]. Taxonomically-relevant DNA methylation under controlled conditions corresponded to chromosomal features and segregating *cis*-acting genetic variation between species, tempering expectations for an independent role of epigenetic variation during nascent species divergence [24]. For a small subset of sites, the study further provides evidence for environmentally induced methylation variation, though with unclear intra-(soma-to-soma) and transgenerational (soma-to-germline) heritability or functional relevance. We conclude that the main source of methylation variation in natural populations of crow is predetermined by chromosomal organization and genetic variation and provides little scope for independent evolution at this early stage of species diversification. Whether this finding is limited to animals where the possibility of autonomous epigenetic inheritance hinges on the degree of residual epigenetic variation surviving deterministic reprogramming [16,116] motivates further study.

## Notes

### Ethical approval

All applicable international, national, and/or institutional guidelines for the care and use of animals were followed. Permission for sampling of wild carrion crow was granted by *Regierungspräsidium Freiburg* (Aktenzeichen 55-8852.15/05), 8852.15), Landratsamt Zwickau (364.622-N-Her-1/14), Landratsamt Mittelsachsen (55410704 Beringungserl-Voigt_14), Lanjdratsamt Vogtlandkreis (364.622-2-2-88841/2014), Landratsamt Meißen (672/364.621-Kennzeichnung von Tieren 18935/2013), Landratsamt Bauzen (67.3-364.622:13-01-Krähen), Landesdirektion Sachsen (24-9168.00/2013-4), Landesamt für Verbraucherschutz, Landwirtschaftund Flurneuordnung Brandenburg (23-2347-8a182008) in Germany and by Jordbruksverket (Dnr 30-1326/10) in Sweden. Import into Sweden was registered with *Veterinäramt Konstanz* (Bescheinigungsnummer INTRA.DE.2014.0047502) and *Jordbruksverket* (Diarienummer 6.6.18-3037/14). Animal husbandry and experimentation was authorized and inspected on site by *Jordbruksverket* (Diarienummer 5.2.18-3065/13, Diarienummer 27–14) and ethically approved by the European Research Council (ERCStG-336536). Crows were hand-raised and thereafter tightly monitored by experienced staff including one animal caretaker present throughout the entire period of 2 years. The social behaviour of the group was monitored on several occasions daily, from close-by during feeding and from outside the aviary. Health status was monitored on regular intervals by staff and external veterinaries. While aggressive behaviours over food access (e.g. threatening) or perching sites (e.g. displacement) could be observed, none of the crows showed aberrant behaviour or a visible indication of stress.

## Acknowledgments

The sampling for this study would have been impossible without the help of ornithologists who identified active nest sites and assisted in sampling. Special thanks to M. Hug and colleagues in Brandenburg, J. Voigt, D. Kronbach, J. Wollmerstädt, M. Schrack and W. Nachtigall in Sachsen, M. Döpfner in Baden-Württemberg, S. Zinko and M. Grossmann in Austria, and personnel of the Amministrazione Provinciale of Alessandria, Asti and Cuneo in Italy. The Max Planck Institute for Ornithology in Radolfzell, the Friedrich-Löffler-Institute and the Förderverein Sächsische Vogelschutzwarte Neschwitz e. V. further assisted with the organization of field work. Computing was performed on the BioHPC hosted at Leibniz Rechenzentrum Munich funded by the German Research Foundation. The authors also acknowledge Alice Merondun for the original crow illustrations used throughout the manuscript.

## Funding

European Research Council grant ERCStG-336536 FuncSpecGen (JBWW)

Knut and Alice Wallenberg Foundation project grant (JBWW)

LMU Munich (JBWW)

German Research Foundation grant WO 1426/2-1 (JBWW)

German Research Foundation grant INST 86/2050-1 FUGG (JBWW)

## Author Contributions

Conceptualization: JM, JBWW

Methodology: JM, JBWW

Investigation: JM, JBWW

Visualization: JM, JBWW

Funding acquisition: JBWW

Project administration: JBWW

Supervision: JBWW

Writing – original draft: JM

Writing – review & editing: JM, JBWW

## Data and Materials availability

All associated sequence data are uploaded to the NCBI SRA with run accessions indicated in Supplementary **Table S**1 under Bioproject PRJNA594256.

All bioinformatic code and input files to reproduce the figures and tables within the manuscript are available at this GitHub repository https://github.com/merondun/corvus_5mC_divergence, with large input files stored at the manuscript’s Zenodo repository: https://zenodo.org/doi/10.5281/zenodo.7641838.

## Supporting Information for

### The contribution of epigenetic variation to evolution in crows

Justin Merondun, Jochen B. W. Wolf

### Supplementary Information Includes

Supplementary Text

Figs. S1 to S24

Table Captions S1 to S15

### Other Supplementary Materials include

Tables S1-S15 (Tables.xlsx)

External Repository with Bioinformatic code: https://zenodo.org/doi/10.5281/zenodo.7641838

### Supplementary Text

#### Genome properties and *cis*-genetic variation

We examined broad-scale relationships between DNA methylation divergence and its chromosomal substrate (population genetic variation, genomic annotation features, chromosome length, positioning along a chromosome, GC-content) primarily using a supervised machine learning regression approach. We complemented this approach with a supervised machine learning classification approach to see if models could capture more variation when only classifying sites into ‘Low’ and ‘High’ levels of divergence. Our reasoning for this approach was to see if discretization of our response variable (DNA methylation divergence) increased the relationships to genome properties. For instance, if DNA methylation divergence operated differently with genome properties at levels of ‘High’ and ‘Low Divergence’ instead of strictly along a continuum of divergence values, perhaps the relationship with explanatory variables would be stronger with a binary approach. We also extended this approach into a trinary response variable, corresponding to ‘Low’, ‘Intermediate’, and ‘High’ levels of DNA methylation divergence. Our DNA methylation divergence response variable, the binned DMP test -statistic estimates for taxon effects in 5-Kb windows, was transformed into a binary or trinary variable corresponding to quantile distributions within each experiment. For instance, within the ComGar.WGBS experiment to create a binary DNA methylation response, we set all methylation divergence values above the 80% quantile to ‘High’, and all windows less than the 80% quantile to ‘Low’. Similarly, for a trinary response we set a lower bound at 20%, so windows falling between the 20 – 80% quantile in the distribution are classified as ‘Intermediate’. This was repeated for each experiment, corresponding to the distributions and thresholds detailed in **<u>Fig</u> <u>S15</u>**. Similar covariate permutation importance values were recovered with this approach (**<u>Fig</u> <u>S13</u>**).

### Supplementary Table Captions (*see .xlsx)*

**<u>Table S1.</u> Metadata for the four experimental set-ups (ComGar.RRBS, ComGar.WGBS, HybZon.RRBS, Whole-genome resequencing)**. Genotype and Hybrid indices were measured in an external study (Knief et al. 2019). Weather data was obtained from Meteostat for the closest locality for the 3 months preceding hatch date, with a link provided.

**<u>Table S2.</u> Aligned and deduplicated read pairs and median genome-wide coverage for the 28 male individuals from the resequencing reanalysis.**

**<u>Table S3.</u> Total read counts, mapping efficiency, and bisulfite-conversion efficiency across the three DNA methylation experiments.**

**<u>Table S4.</u> Determinants of global methylation variation.** Significance of physiological or environmental variables for explaining global patterns in DNA methylation across all three experiments. First, the optimal distance metric was chosen based on Spearman correlations between a data frame of methylation states and explanatory variables using function ‘rankindex’ in R. Constrained ordinations using vegan were estimated using the supplied distance, with significance of individual variables (or terms) assessed with 10,000 permutations using the ‘anova’ function. An analysis of similarity was also performed on the NMDS output for taxon and environmental variables, also using 10,000 permutations.

**<u>Table S5.</u> Four taxon-related dual-experimentally validated DMPs.** Details on the 4 taxon-related DMPs, identified as significant with a chromosome 18 genetic hybrid index and identified as ‘taxon’ related in the common garden experiment, and the 3 pre-hatch temperature related DMPs, identified as conserved in the common garden experiment, and significant with temperature in the hybrid zone experiment (cf. Fig 1, Fig 2).

**<u>Table S6.</u> Overlap between each DNA methylation experiment and the population whole-genome resequencing dataset.** Genetic windows were calculated in 5-Kb genome-wide, and have lower overlap with the RRBS datasets, seemingly because of missing resequencing data in areas of high GC content.

**<u>Table S7.</u> Pairwise correlations between explanatory variables used for supervised machine learning.** Relationships between taxonomic DNA methylation divergence (test-statistic estimates from DMP analyses) were averaged in 5-Kb windows (the resolution of the population genetic data), and were modeled with random forests and boosted regression trees using a regression approach.

**<u>Table S8.</u> Machine learning model fit.** The entire modelling procedure was replicated 3 times providing the error estimates. The process was repeated both as a classification problem with binary and trinary responses (binned by quantile thresholds), or as a regression problem, using both random forests and boosted trees. The response variable (taxonomic-specific test-statistic from beta-binomial regressions using DSS) was computed within 5-Kb windows either as the mean value (Mean) or the maximum value (Max).

**<u>Table S9.</u> Machine learning covariate permutation importance.** The entire modelling procedure was replicated 3 times providing the error estimates. The process was repeated both as a classification problem with binary and trinary responses (binned by quantile thresholds), or as a regression problem, using both random forests and boosted trees. The response variable (taxonomic-specific test-statistic estimates from beta-binomial regressions using DSS, cf. Fig 4) was computed within 5-Kb windows either as the mean value (Mean) or the maximum value (Max).

**<u>Table S10.</u> Differentially methylated points (DMPs) according to each physiological or environmental classification.** Single experimental classification represents the total DMPs if only a single experimental set-up was used to detect DMPs. Multi-experimental validation required at least 2 corroborative classifications from the 3 experiments to identify a site (cf. Fig 2).

**<u>Table S11.</u> Missingness between experiments after base-pair level CpG merging, used for DMP classifications.**

**<u>Table S12.</u> Counts of total differentially methylated point (DMP) classifications.** Final classifications are determined via the strategy outlined in Figure 2. Missingness: Whether or not the CpGs were present in all experiments, 2 experiments, or 1 experiment. The remaining columns are binary (0/1) descriptors indicating whether or not the condition of that column is satisfied. For necessary and sufficient conditions, refer Fig. 2.

**<u>Table S13.</u> Population genetic metrics within and outside of the focal region on chromosome 18 (start 8.07e6; end 10.07e6), harboring 100% of the SNPs with F_ST_ > 0.5, and 99% of the SNPs with F_ST_ > 0.3.** Empirical values for mean, minimum, and maximum from the genome-wide data are also shown below.

**<u>Table S14.</u> Mean taxonomically-related DNA methylation divergence (absolute value of taxon test statistic estimates from beta-binomial regressions) within and outside of the focal region on chromosome 18.** Due to unequal sample size we sampled the number of focal windows present within each comparison (indicated below), and sampled those number of windows within the focal region and in the autosomal background 1,000 times with replacement. If either side of the 95% CI distributions overlapped, they were not considered significantly different. DNA methylation divergence was assessed at base-pair level.

**<u>Table S15.</u> Spearman’s rank correlations between population genetic metrics and mean taxonomically-related DNA methylation divergence (absolute value of test statistic from beta-binomial regressions) within and outside of the focal region on chromosome 18.** Lower and upper 95% quantiles drawn from 1,000 bootstrap replicates. Test statistics came from DMP detection analyses, and were averaged (mean) in 5-Kb windows, in which the genetic data was analyzed. We observed qualitative higher correlations between DNA methylation divergence and population genetic metrics within the focal island, but only D_XY_ and haplotype diversity had 95% CI distributions that did not overlap 0 within the focal region (cf. Fig 4).

### Supplementary Figures

**Fig S1.**
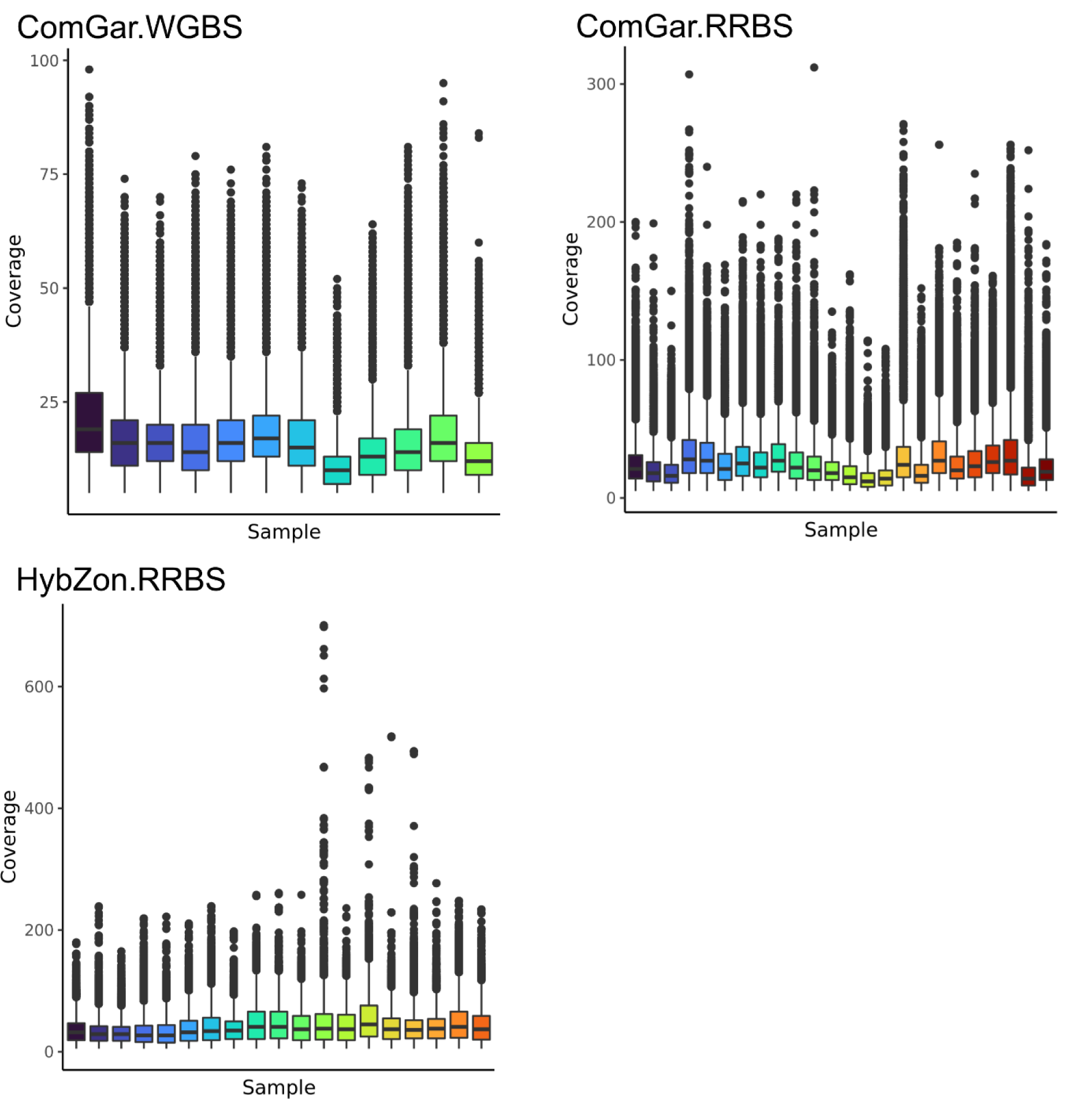
DNA methylation sequencing coverage. Sequencing coverage from the three bisulfite sequencing experiments. Base-pair resolution counts for CpG sites are summarized for all samples (x-axis), with default parameter boxplots showing variation across each experiment. Visualized with R.

**Fig S2.**
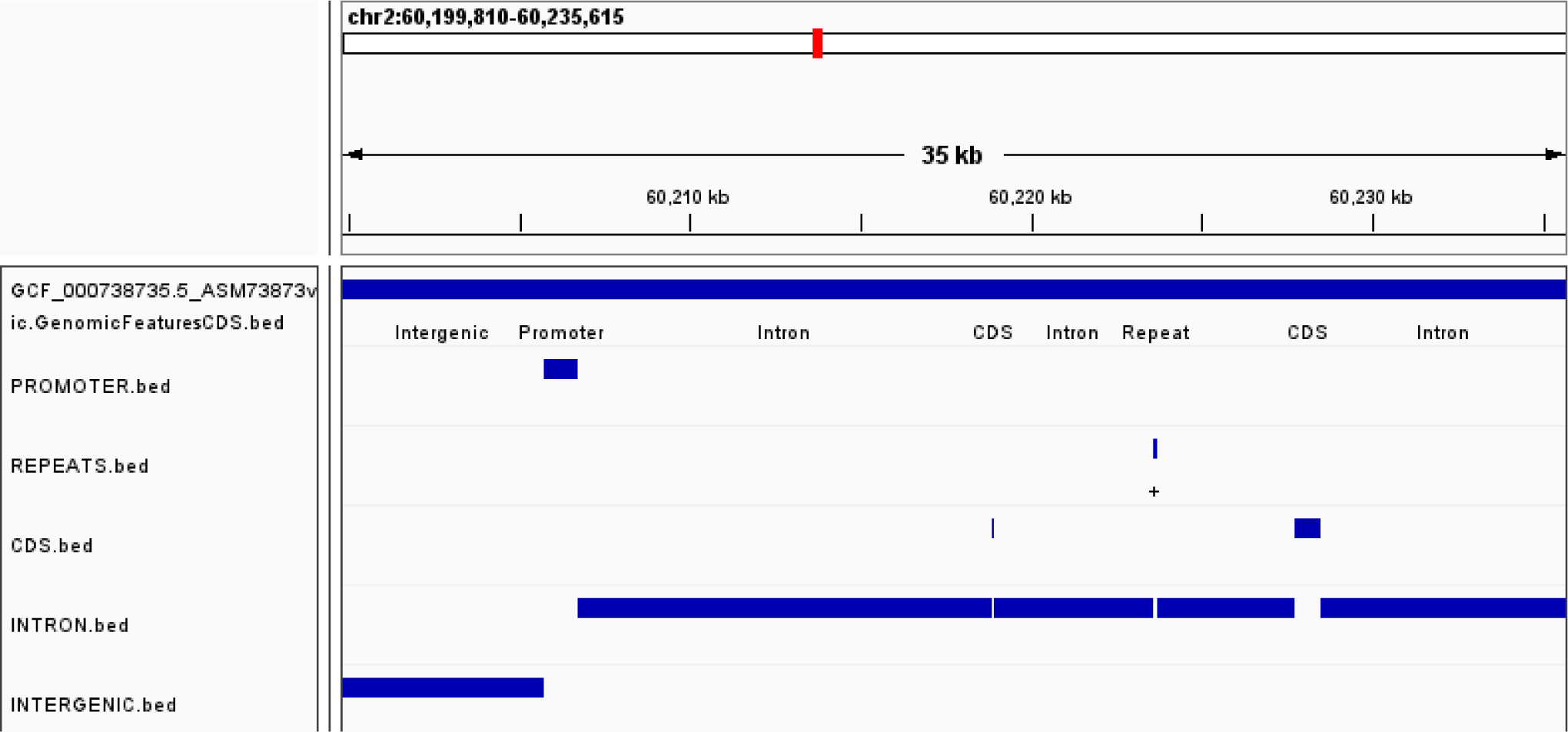
Genomic feature designation. Example Annotation Feature track for the European crow reference genome. Promoters were identified as CpG islands that overlap the 2-Kb region upstream of a gene start. Repeats were identified from RepeatMasker (excluding simple repeats). Genic coding sequencing (CDS) and intronic regions were taken directly from the RefSeq annotation. Remaining regions were annotated as intergenic. Sites were assigned a single genomic context, with priority given in the order as indicated: promoters, repeats, CDS, intron, intergenic. Visualized with the integrative genome browser.

**Fig S3.**
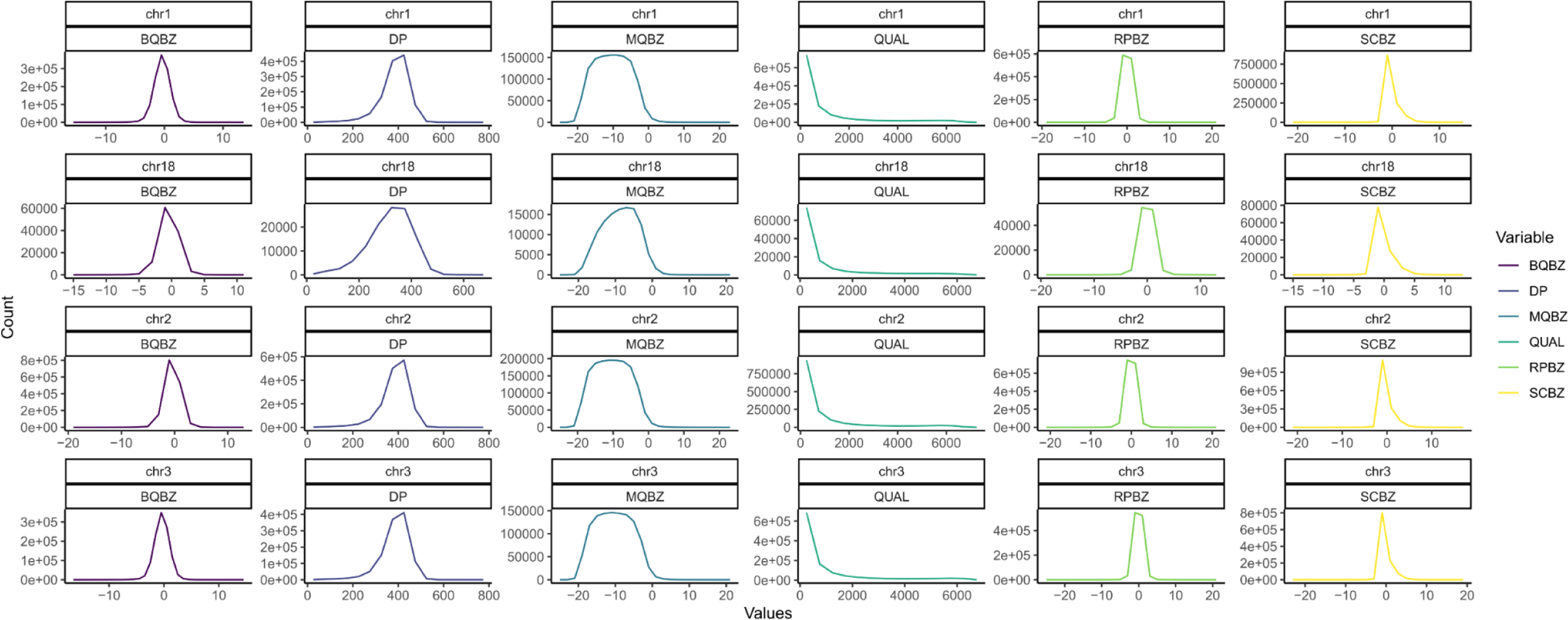
Whole genome resequencing SNP quality metrics. Resequencing experiment SNP quality, showing 6 INFO metrics for the 28 hooded and carrion crows used to assess C-T and A-G SNPs and quantify genetic variation. We applied hard depth and quality filters to filter SNPs (*QUAL > 20, DP < 2*Average DP*, or *DP > 28x*). Only 4 chromosomes are shown for brevity. BQBZ = base quality bias; DP = depth; MQBZ = map quality bias; QUAL = quality; RPBZ = read position bias; SCBZ = soft-clip length bias. Visualized in *R* with the *tidyverse*.

**Fig S4.**
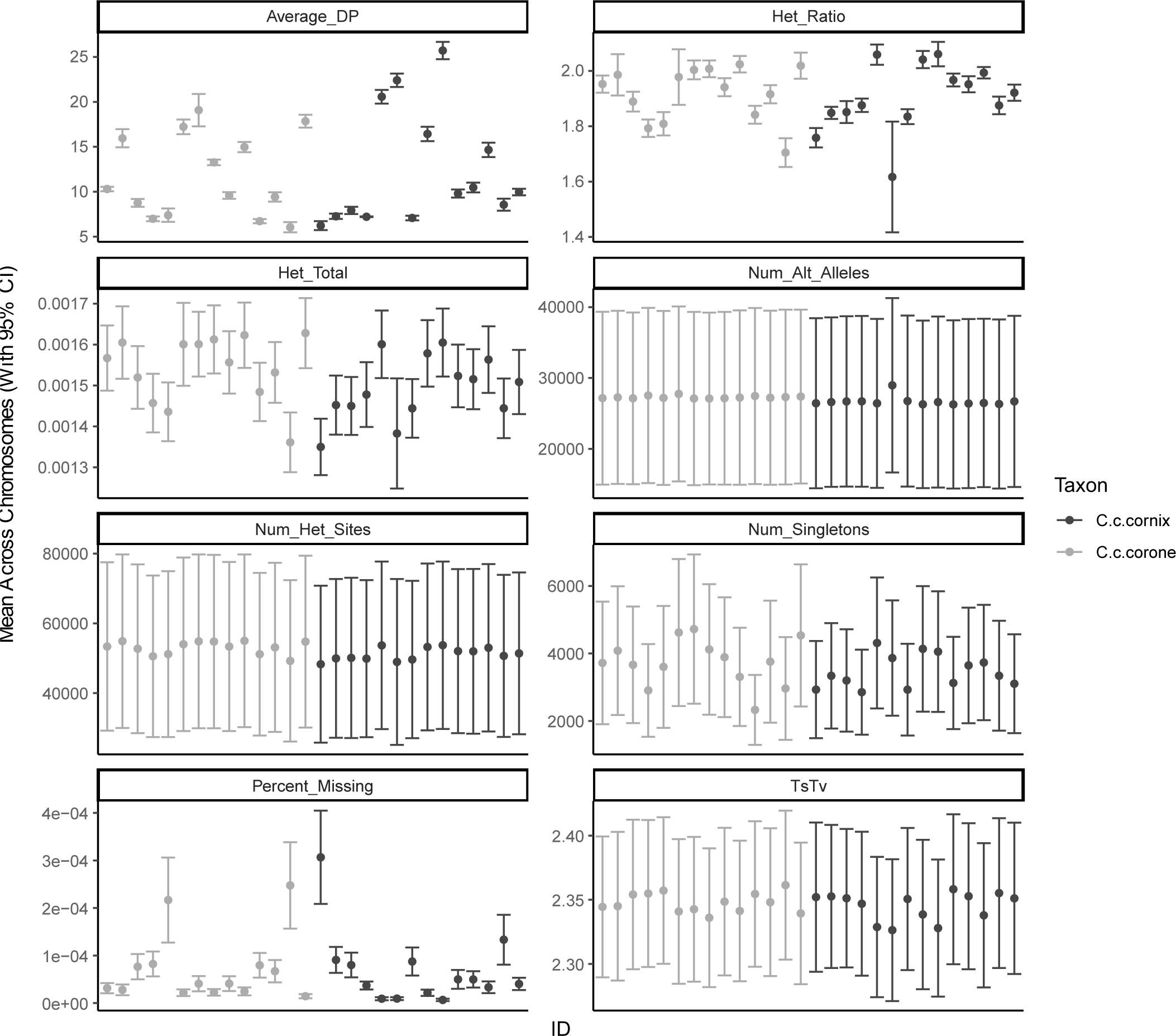
Whole genome resequencing SNP genotype quality metrics. Resequencing experiment SNP genotype summaries, showing individuals along the X-axis and values along the Y-axis for 8 different metrics. Means and 95% confidence intervals calculated from the averages of all chromosomes. Average_DP = average depth; Het_Ratio = heterozygosity ratio; Het_Total = total observed heterozygosity; Num_Alt_Alleles = number of alternate (non-reference) alleles; Num_Het_Sites = number of heterozygous sites; Num_Singletons = number of singletons; Percent_Missing = percentage of missing genotypes; TsTV = transition to transversion ratio. Visualized in *R* with the *tidyverse*.

**Fig S5.**
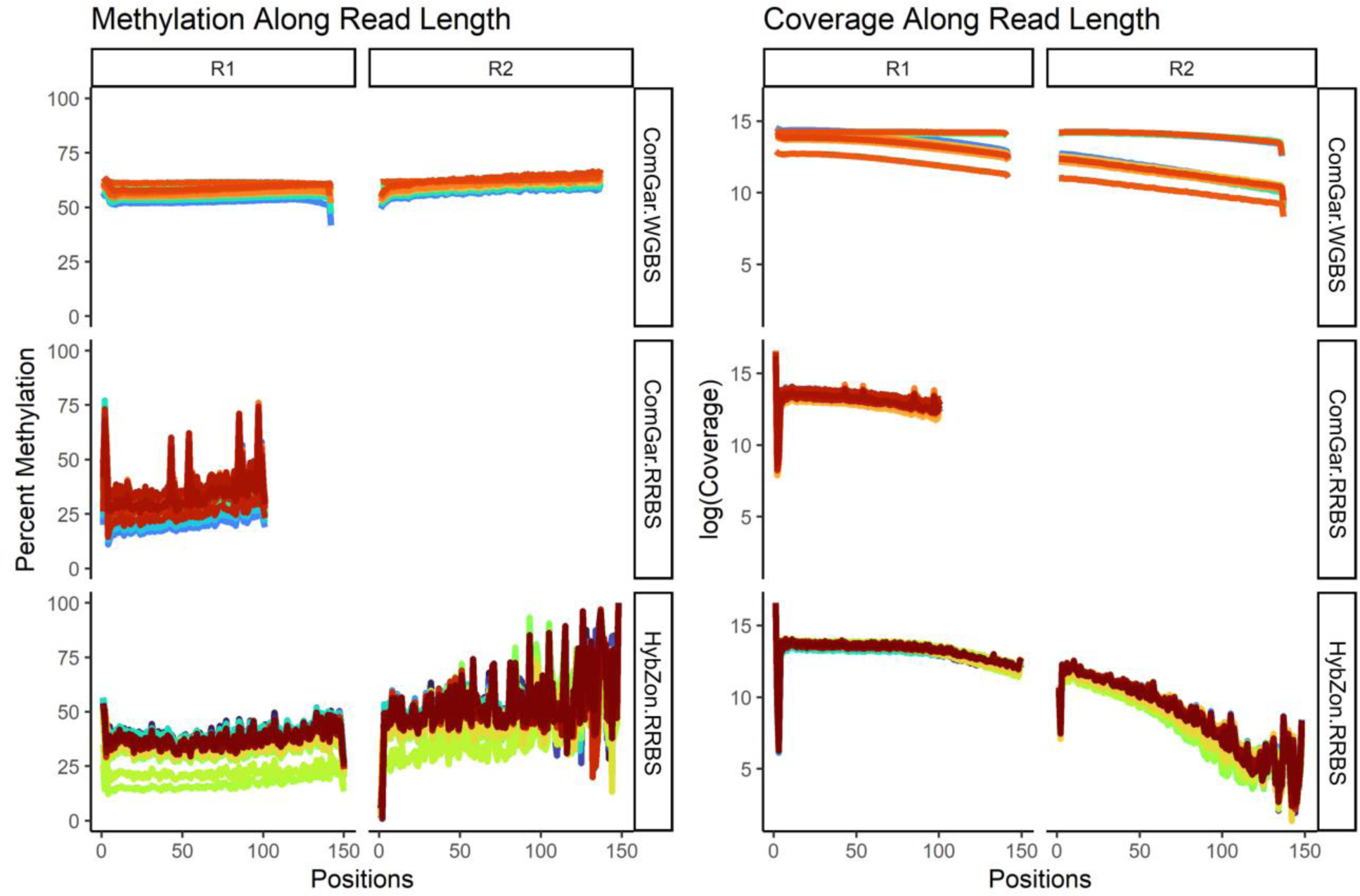
Methylation-bias and coverage along read length for all libraries. Methylation bias plots showing mean methylation levels and total (log-transformed) coverage across read lengths from the three bisulfite sequencing experiments, with each individual color-coded. Nine bases were trimmed from the R1 and R2 5’ ends of the WGBS reads to control for the post-bisulfite adapter tagging artifacts, as well as 1 base from the 3’ end. RRBS reads were trimmed with the ‘—rrbs’ flag of trim_galore. The spikes in RRBS reads appear to arise from high coverage outliers, which are removed in the analysis. The three individuals in the HybZon.RRBS experiment which exhibit systematic hypomethylation were further analyzed and removed from the analysis due to potential external confounding biological factors. Visualized in *R* with the *tidyverse*.

**Fig S6.**
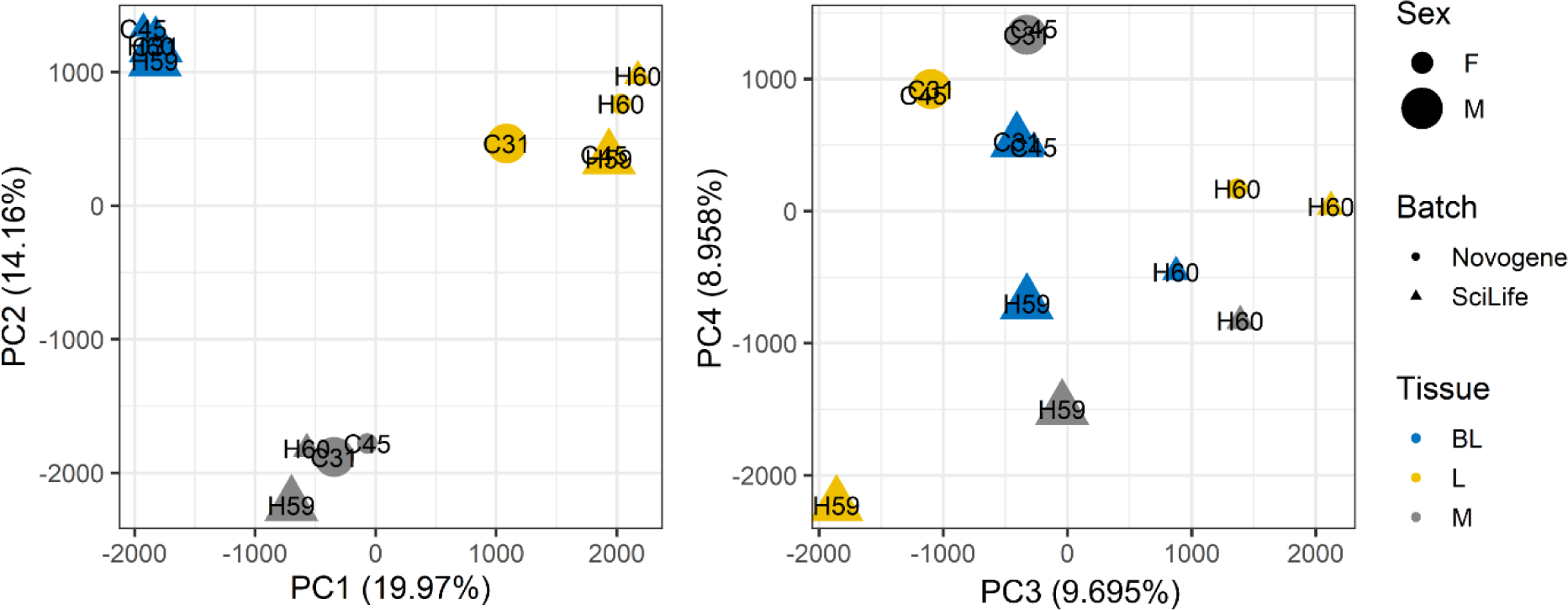
Batch effects for ComGar.WGBS libraries. Assessing batch effects from the two sequencing centers (Novogene, SciLife) for the ComGar.WGBS dataset, with sexes denoted with size, tissues by color, and shape by sequencing center. Based on the first four axes of a scaled and centered PCA ordination on total DNA methylation data, no batch effects warranted finer scale analysis. Ordinations completed with *prcomp* in *R*.

**Fig S7.**
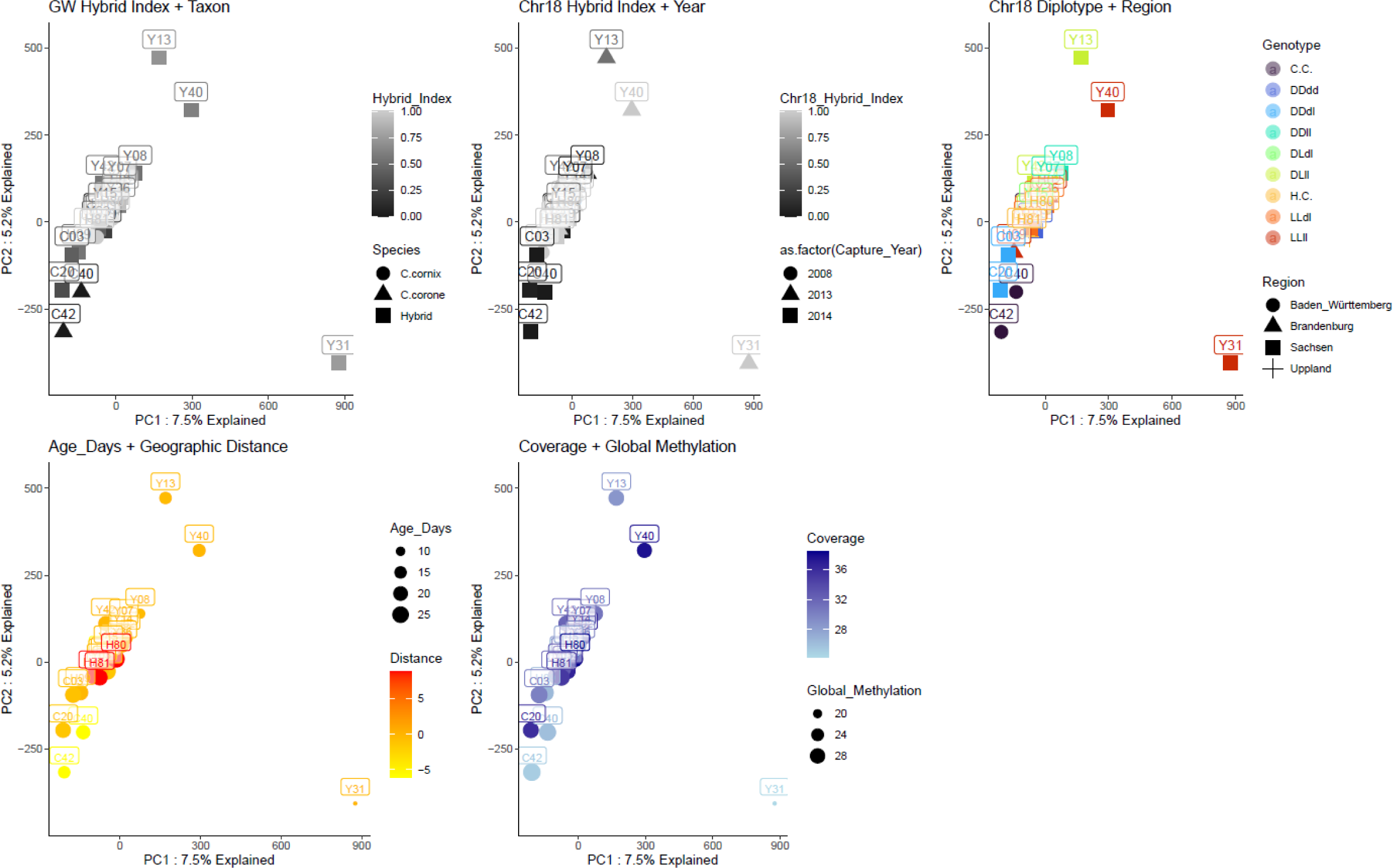
HybZon.RRBS outlier individual quantification. Three individuals (Y13, Y40, Y31) in the HybZon.RRBS experiment appeared as outliers in PCA ordinations of all DNA methylation sites. The PCA ordinations above show different symbology according to potential confounding factors driving these samples as outliers. These samples appear to exhibit some systemic hypomethylation and low coverage and were dropped from subsequent analyses.

**Fig S8.**
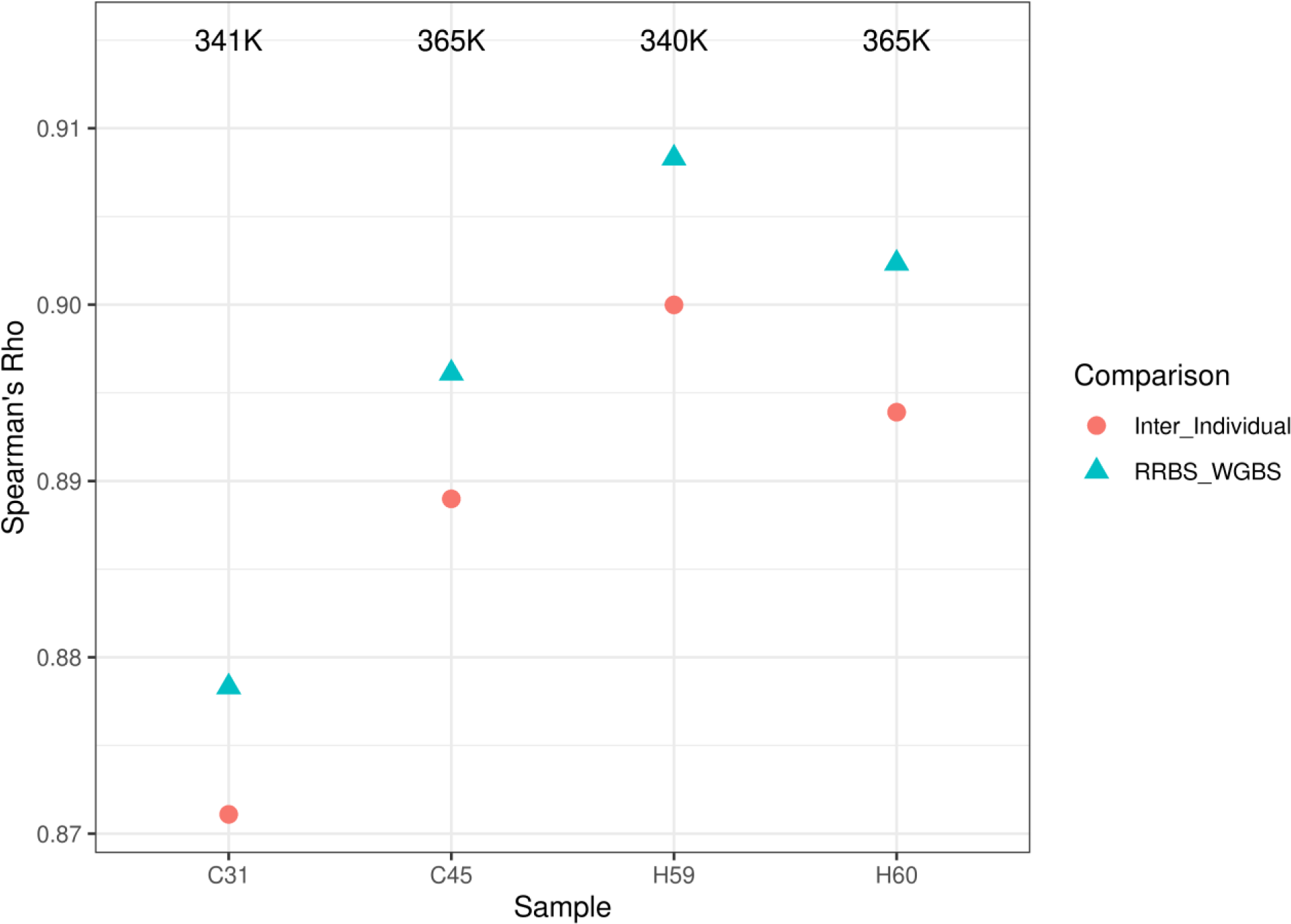
Correlations of DNA methylation between and within individuals. **Spearman’s** rank correlations between RRBS and WGBS biological replicates in the ComGar. experiment, with a randomly sampled inter-individual correlation for comparison. Labels indicate the number of CpGs considered for the correlation, where each correlation was filtered first to retain only sites with no missing data between the biological replicate and one randomly sampled inter-individual comparison. Correlations were calculated in *R* and plotted with *ggplot2*.

**Fig S9.**
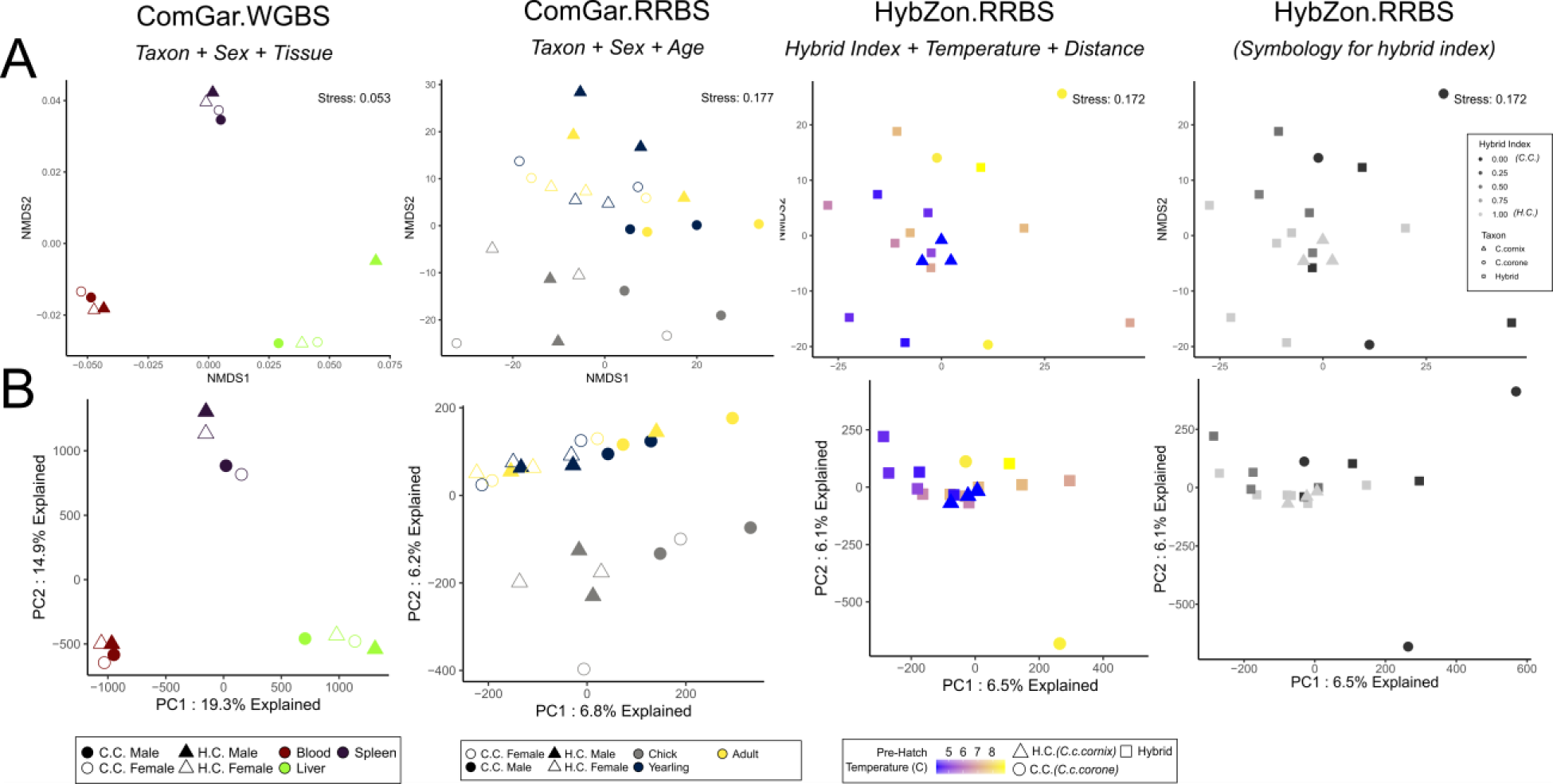
Ordinations for DNA methylation sequencing data across the three experiments. (A) Unconstrained non-metric multidimensional scaling (NMDS) of DNA methylation sequencing data and, (B) principal component analysis ordinations (PCAs). NMDS relies on distances between samples, with the best-fit distance identified with Spearman’s rank index (either Euclidean, Manhattan, Gower’s, Bray-Curtis, or Kulczynski). The best fit measures were Gower’s distance for the ComGar.WGBS experiment, and Euclidean distance for the ComGar.RRBS and HybZon.RRBS experiments. HybZon.RRBS experiment is shown twice for symbology related to temperature or genetic hybrid index. Ordinations were performed with vegan in R with helper functions from tidymodels.

**Fig S10.**
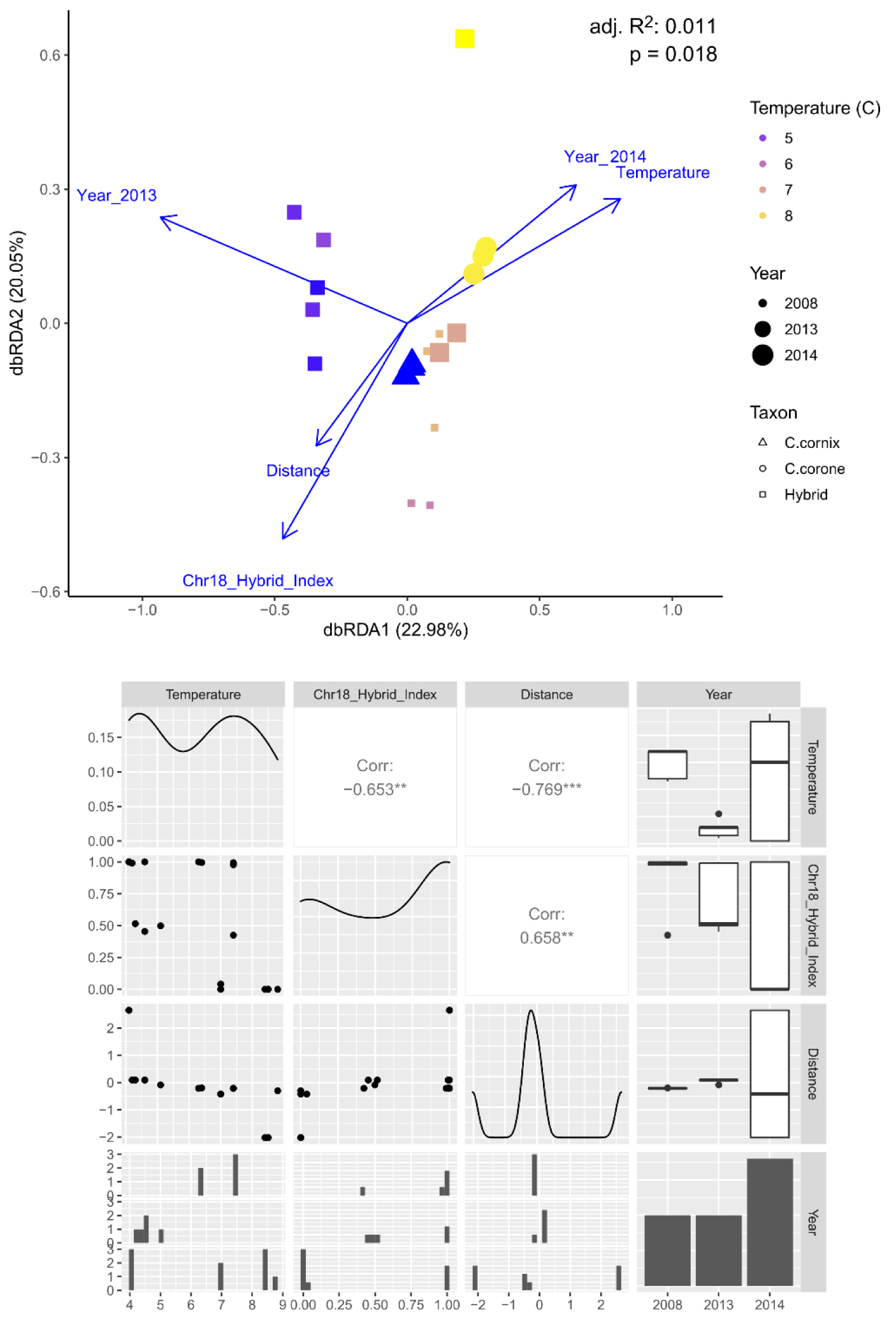
Global determinants of DNA methylation in HybZon.RRBS, using year as an additional covariate. Global associations of DNA methylation variation in the HybZon.RRBS experiment with environmental and genetic covariates, including an additional factor of *Sampling Year* (2008, 2013, 2014). Top panel shows db-RDA biplot with temperature symbolized with color, taxon with shape, and sampling year with size, analyzed and plotted with *vegan* and *ggplot2*. Bottom panel shows Pearson correlations between variables when including the extra factor variable *Sampling Year*, created using *GGally* in *R*.

**Fig S11.**
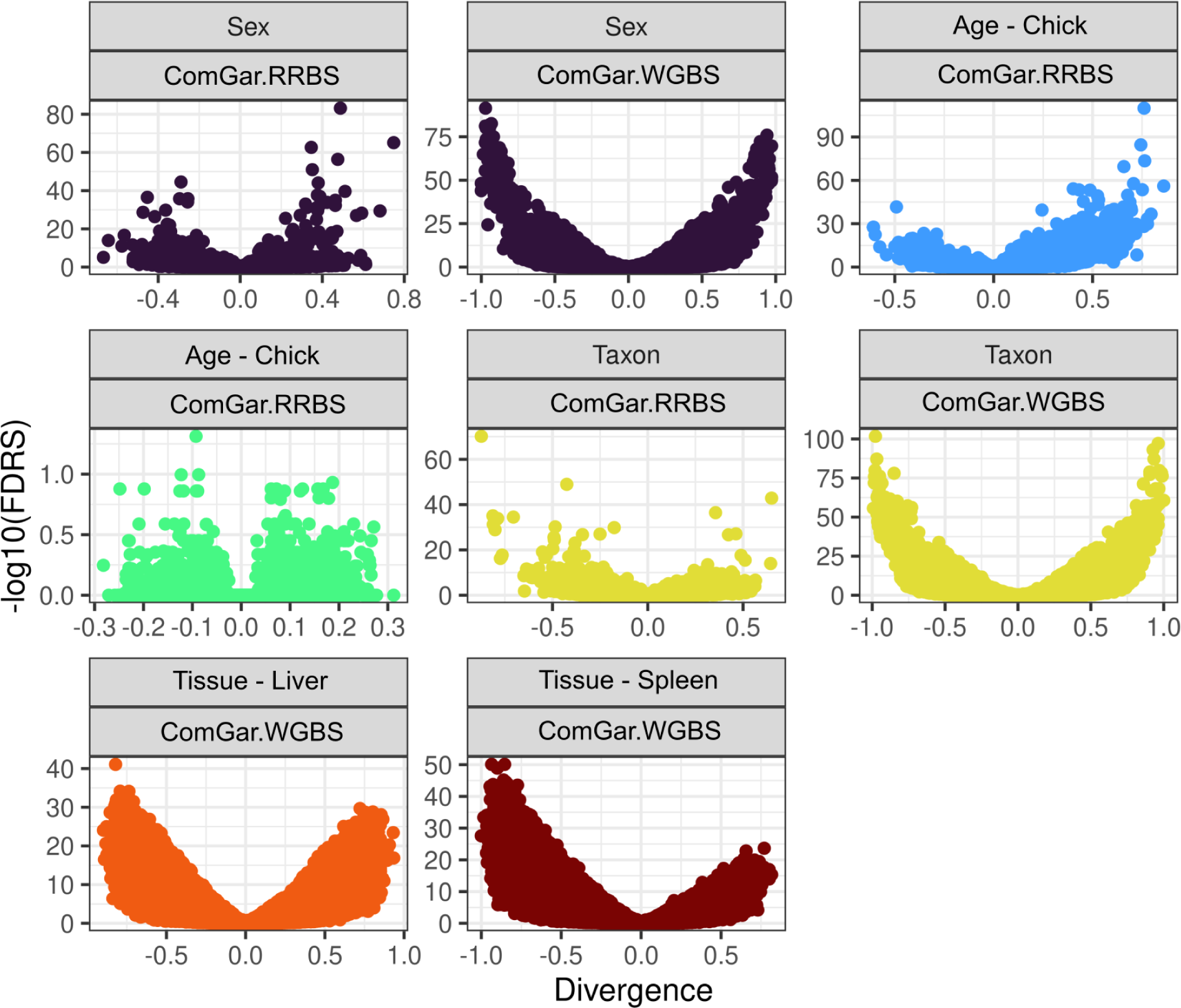
Methylation divergence against DMP significance. DNA methylation FDR-corrected *p*-values from DMP analyses plotted against DNA methylation divergence between groups (ranging from possible −100 to 100% difference). *P-*values come from beta-binomial regressions based on read counts from *DSS* [71]. Divergence values were calculated by subtracting the mean of each group.Visualized in *R*.

**Fig S12.**
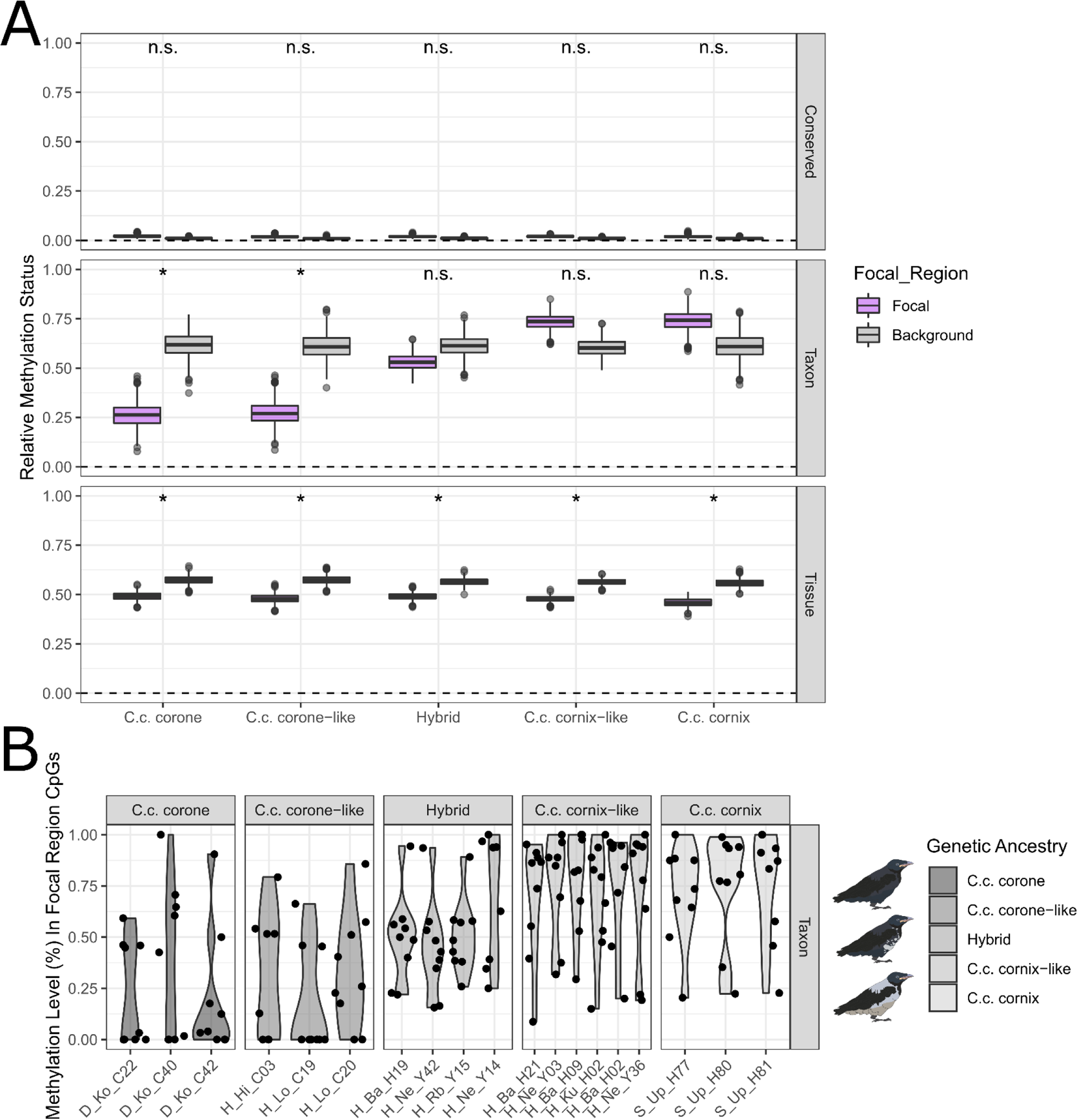
Methylation levels within the HybZon.RRBS experiment, using only classifications from the ComGar. experiments. First, CpG sites from the ComGar.WGBS and ComGar.RRBS were classified into a class (*e.g.,* Taxon, Tissue) based on FDR-corrected p-values from DMP analyses and divergence between groups (see methods). Sites exhibiting less than 10% variation in across the entire experiment are classified as *Conserved*. We then used at least single-experimental validation (*e.g.,* taxon in ComGar.RRBS and either taxon or missing data in ComGar.WGBS) to examine the distributions of methylation levels within the HybZon.RRBS experiment. This is opposed to the multi-experimental validation, which requires significance within the HybZon.RRBS experiment to classify an effect. (**A**) For each class above (*Conserved, Taxon, Tissue*) we show differences in methylation levels within the focal region on chromosome 18 vs the autosomal background with bootstrap sampling. We sampled an equal number of autosomal and focal region CpGs equal to the sites in the focal region with replacement and repeated this1,000 times. Default parameter boxplots show 1^st^ and 3^rd^ quartiles of the bootstrap distributions, significance was indicated if the 95% quartile distributions did not overlap between the focal region and background. Crows within the HybZon.RRBS are divided by hybrid index, ranging from the unadmixed parentals, hybrids with a hybrid index equivalent to either parent, or true hybrids. (**B**) Methylation proportions for each CpG within the focal region for each individual. We observe hypomethylation in *C. c. corone* within the focal region and hypermethylation in *C. c. cornix*, with hybrids intermediate. Visualized with *R* and the *tidyverse*.

**Fig S13.**
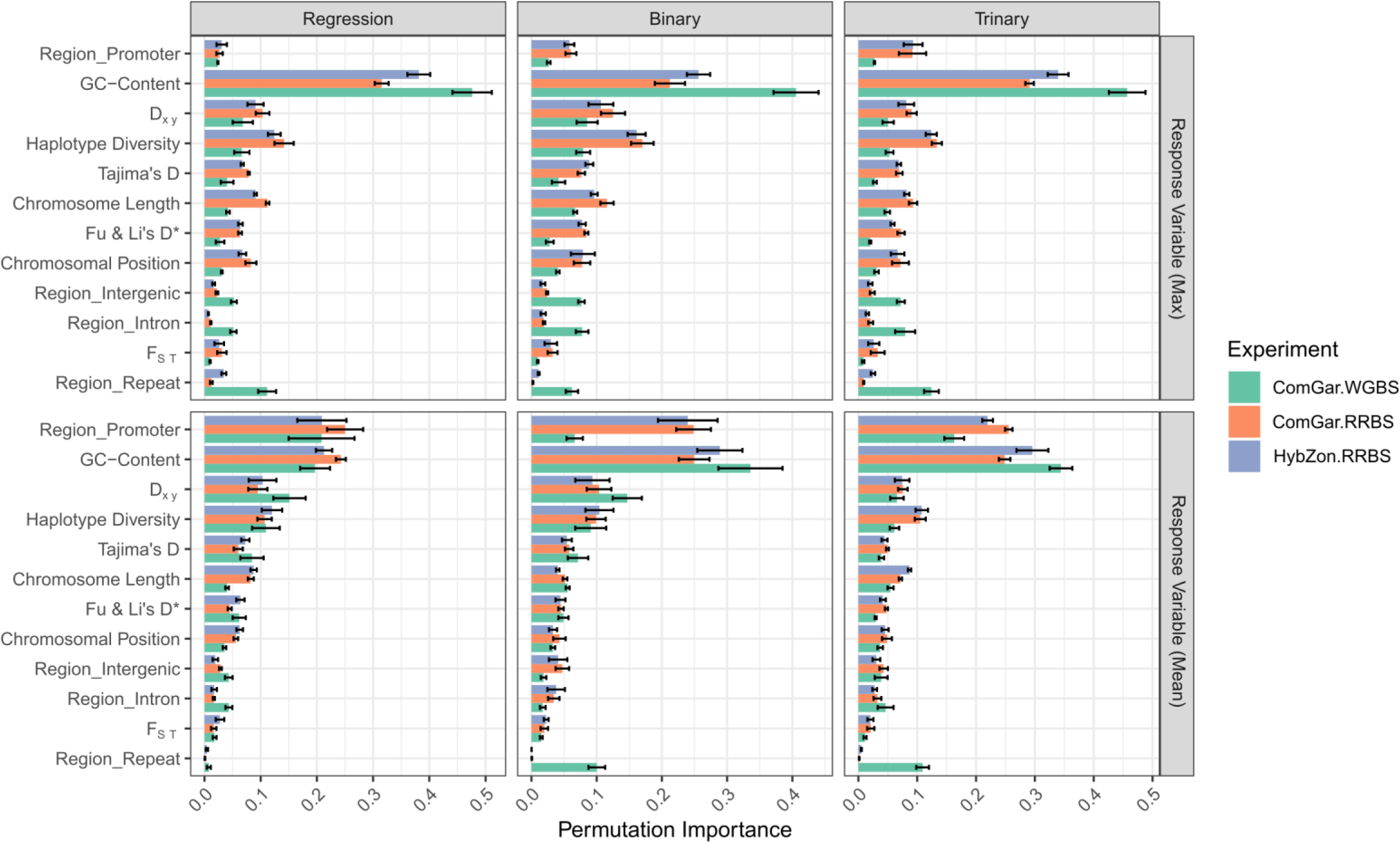
Permutation importance for chromosomal features in determining DNA methylation variation, inferred using machine learning. Covariate permutation importance for all engines and for both mean DNA methylation divergence in 5kb windows, and max DNA methylation divergence in 5kb windows (genetic windows had a resolution of 5kb). To see if large near-zero distributions skewed regression techniques, we also repeated models but with a classification approach using binary and trinary response variables (Low, Intermediate, High; see methods). Cumulative permutation importance was scaled to 1.0 within each iteration to allow comparison between boosted regression trees and random forests. The modelling process was repeated 3 times to generate confidence intervals.

**Fig S14.**
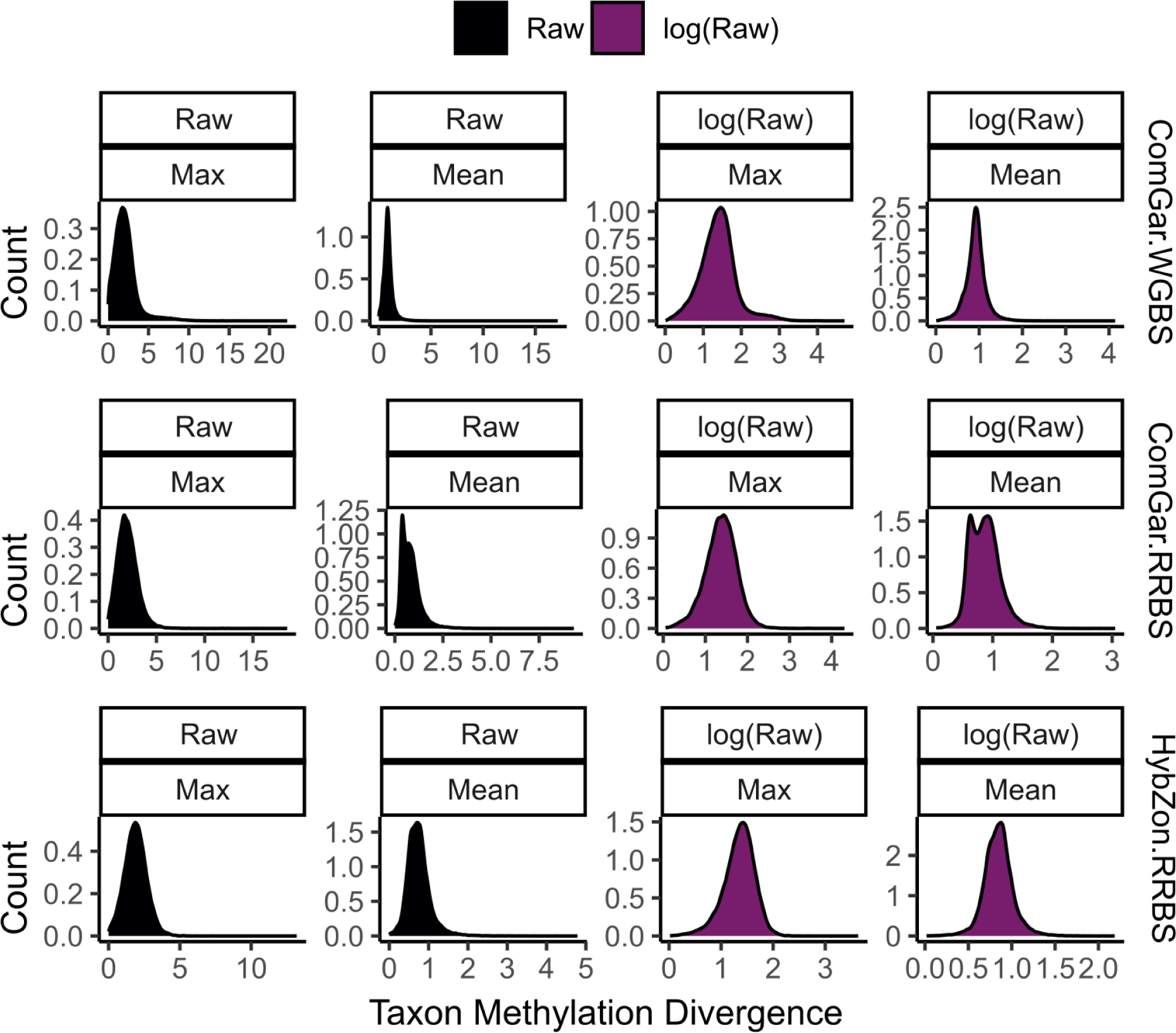
Input response variable distributions for supervised machine learning regressors. These were used for examining the relationship between taxonomic DNA methylation divergence (taxon-specific test-statistic estimates from DMP analyses) and chromosomal determinants (genome property, population genetic variation). Population genetic variation was assessed in 5-Kb windows genome-wide, so the mean or max test-statistics (which were calculated at CpG base-pair resolution) were calculated within each window for sensitivity. We log transformed the response (right panels) prior to modelling. Visualized in *R*.

**Fig S15.**
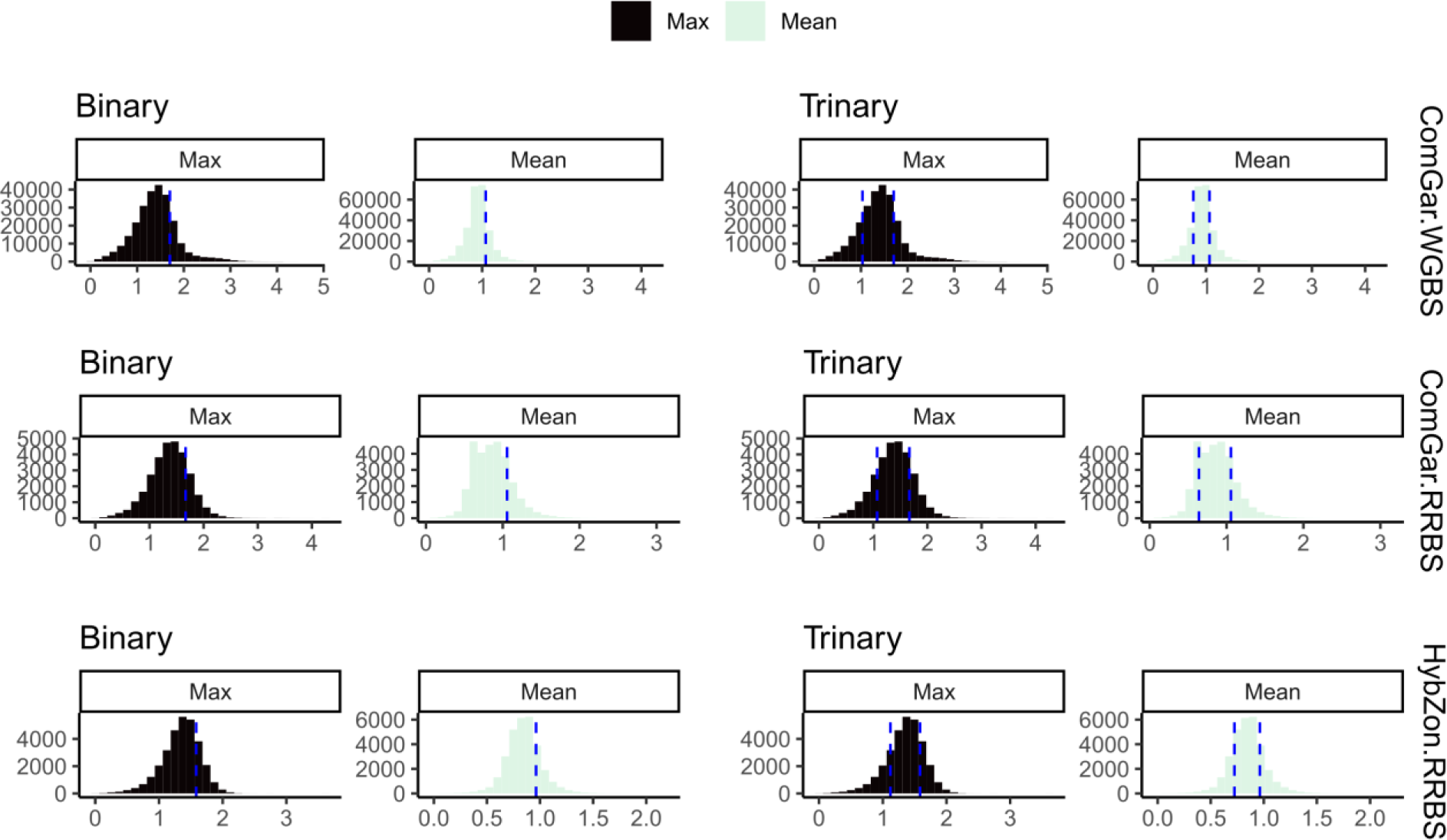
Input response variable distributions for supervised machine learning classifiers. These were used for examining the relationship between taxonomic DNA methylation divergence (taxon-specific test-statistics from DMP analyses) and chromosomal determinants (genome property, population genetic variation). To see if DNA methylation divergence was more parsimoniously modeled as discrete binary or trinary responses, we classified the response variable into binary and trinary categories. Binary categories were assigned as ‘High’ if methylation divergence was above the 80% quantile within each experiment (blue dashed line), and ‘Low’ if below. For trinary responses, we classified ‘High’ above the 80% quantile, ‘Intermediate’ in between 20% and 80%, and ‘Low’ below 20%. We then fit models (random forest, boosted trees), and evaluated models with ROC AUC. Striped lines indicate the threshold above which sites were classified as ‘High’ (binary) or below as ‘Low’, or between which sites are ‘Low’, ‘Intermediate’, and ‘High’ for trinary response. We repeated the analysis except using the maximum taxon-specific estimates within the 5-Kb genomic windows, instead of the mean value. Visualized in *R*.

**Fig S16.**
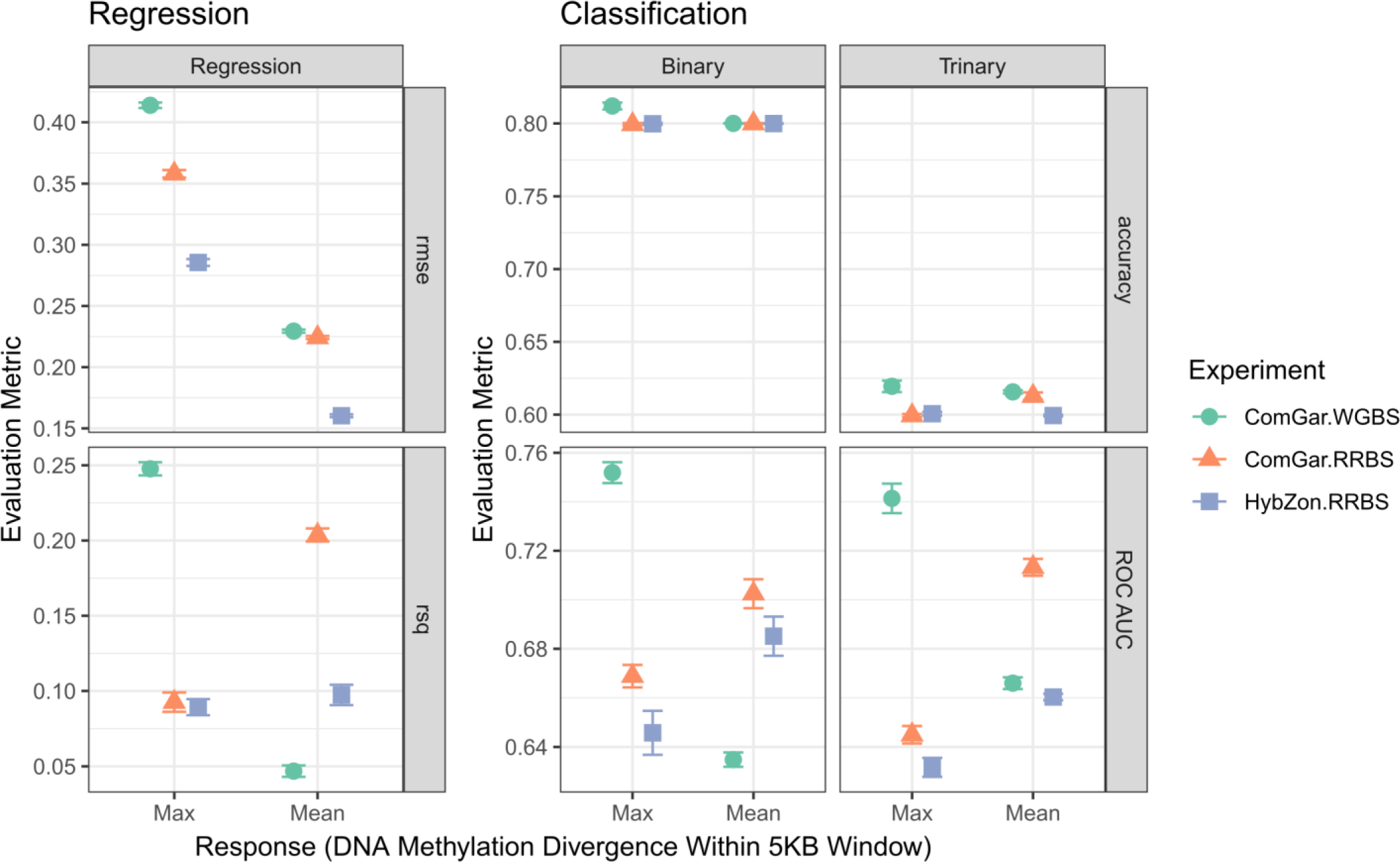
Model fit for regression and classification tasks. Supervised machine learning (random forests, boosted regression trees) were used for assessing the relationship between taxonomic DNA methylation divergence (absolute value of test-statistic estimates from DMP analyses, averaged in 5-Kb windows) and chromosomal features. Values correspond to **<u>Table</u> <u>S8</u>**. Confidence intervals were drawn based on repeating the entire modelling process 3 times. Visualized in *R*.

**Fig S17.**
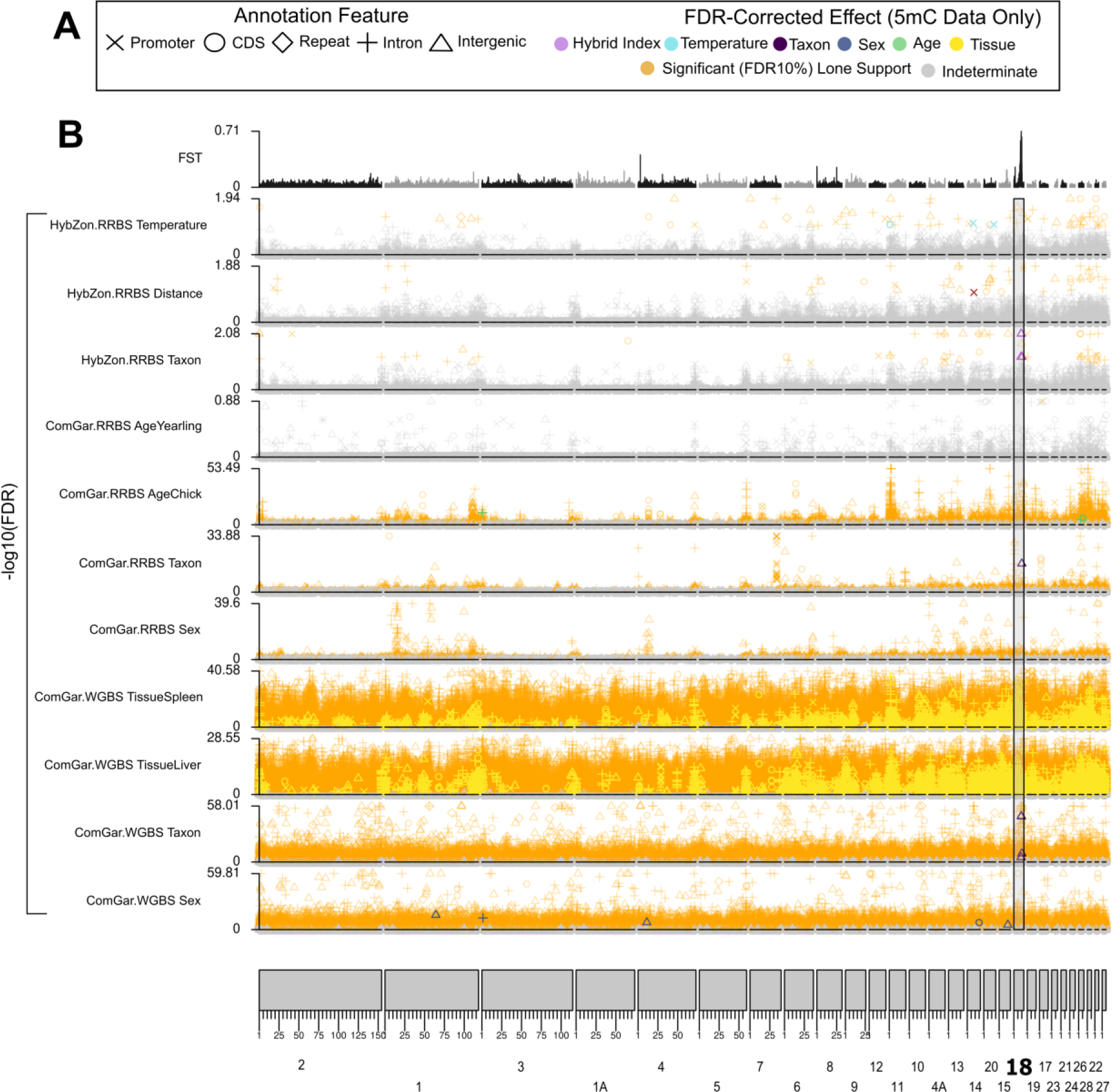
DMPs genome-wide for all variables. DNA methylation (-log10(*p*-value)) and genetic differentiation (F_ST_) data for hooded, carrion, and hybrid crows genome-wide, with a (*A)* common legend. *(B)* CpGs from methylation experiments corresponding to the ComGar.WGBS, ComGar.RRBS, and HybZon.RRBS experiments along the bottom 11 panels showing the FDR-corrected *p*-value for each effect. Points are colored orange if they alone have an FDR < 10%, and grey if they are not significant for the target effect. Points are colored corresponding to the ultimate covariate classification if they have been validated intra-experimentally following the verified classification scheme (***cf.* Fig. 1**). Top track shows genetic differentiation between carrion and hooded crows. Visualized with *karyoploteR*.

**Fig S18.**
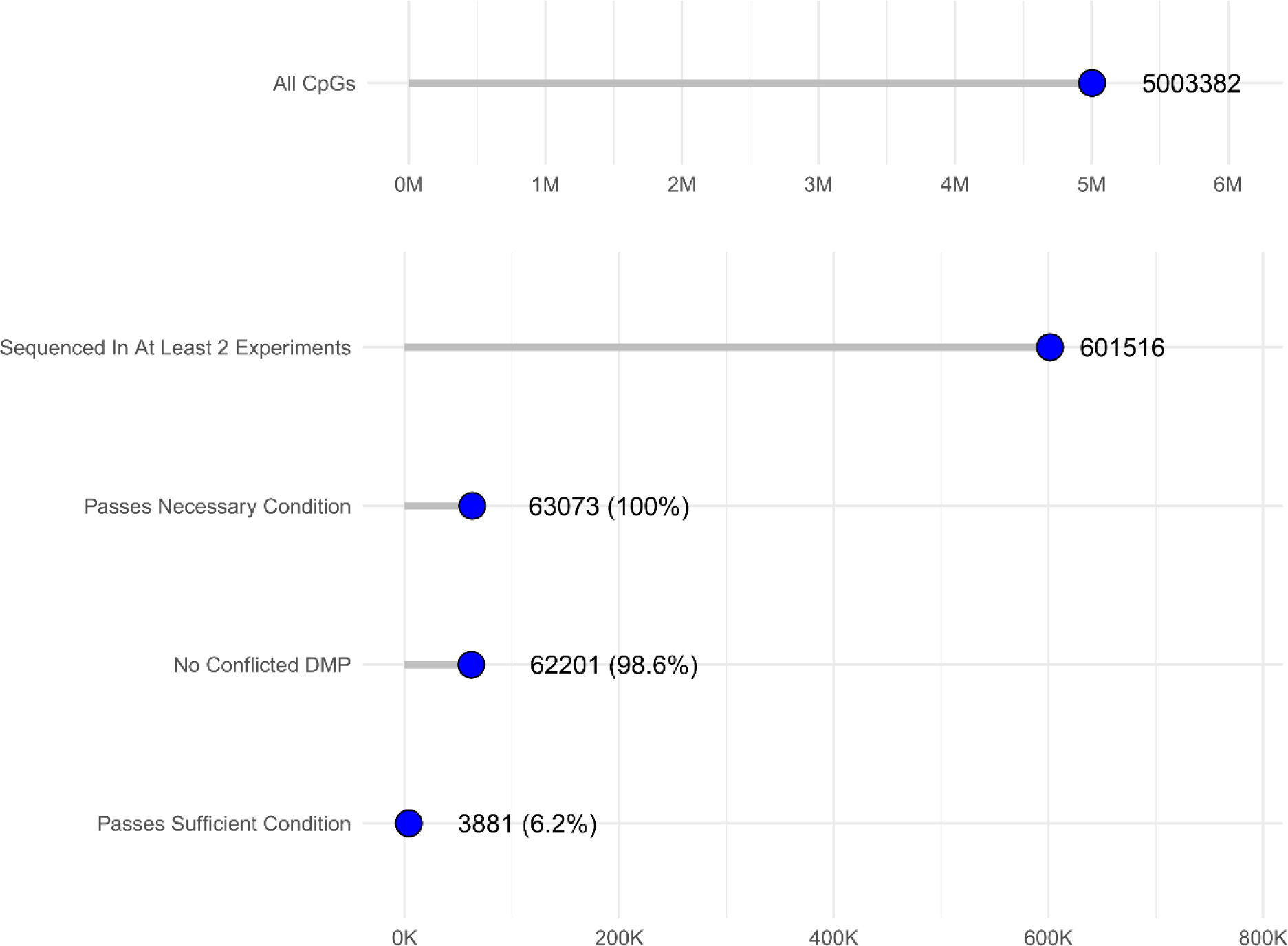
Multi-experimental filtering of CpGs across experiments. Top panel indicates total CpGs sequenced across all 3 experiments. Bottom panel indicates the effects of multi-experimental controls on filtering CpGs. Within the bottom panel, CpGs must have passed the previous filter to be counted for the following filt er (**<u>Table S12</u>**). *DMP For Effect*: after filtering for CpGs sufficiently sequenced in at least 2 experiments (> 10x coverage for RRBS; > 5x for WGBS), these are CpGs that would be classified as DMPs following a traditional single-experimental approach. DMPs were classified based on significance from beta-binomial regressions on methylation counts, and minimum percent difference between groups. *No Conflicted DMP*: DMPs that do not exhibit conflicting DMP designations. *Passes Necessary / Sufficient Conditions*: DMPs that pass their respective conditions ().

**Fig S19.**
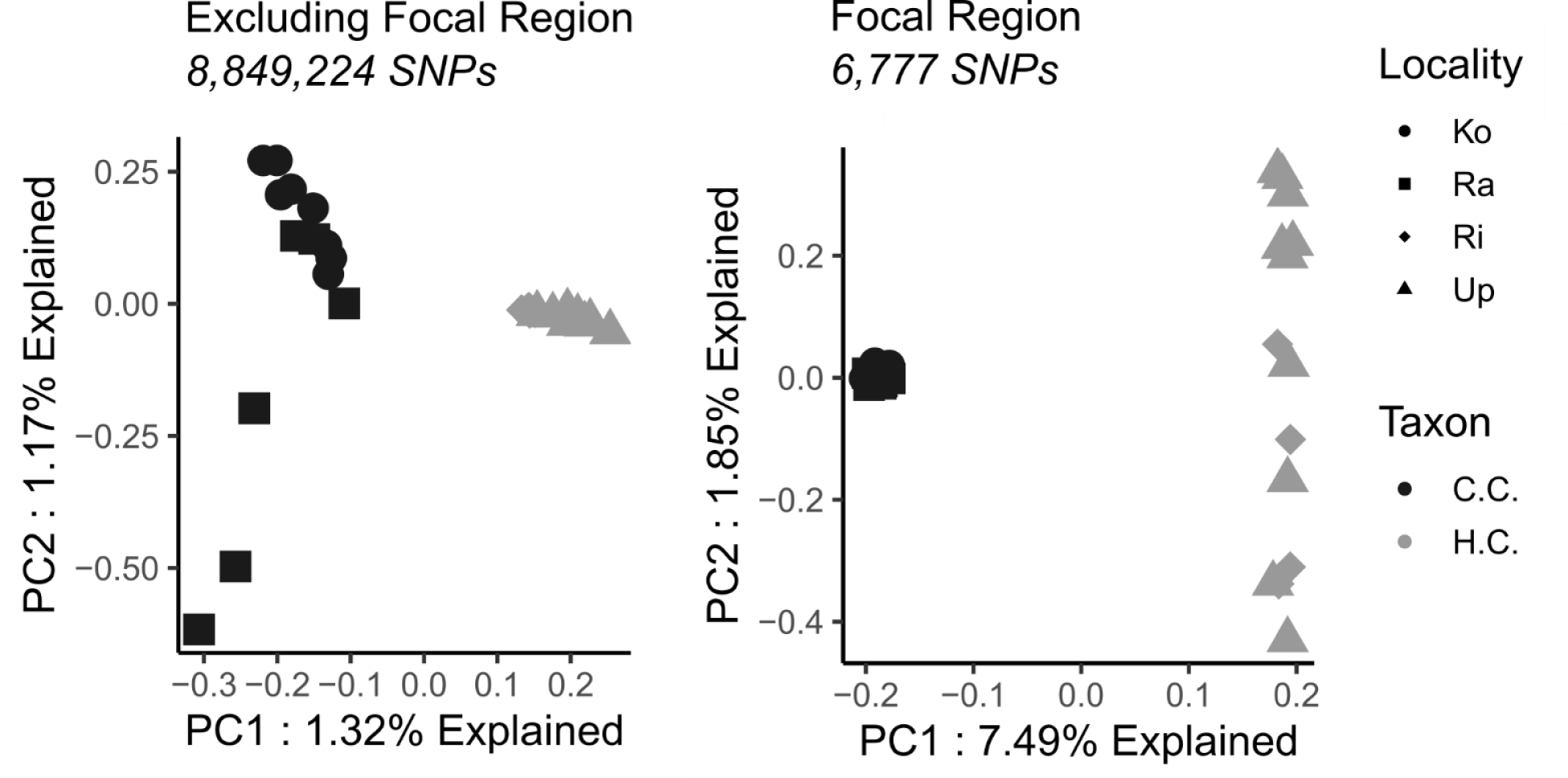
Principal components analysis on genetic whole-genome resequencing data on autosomal SNPs excluding the focal region on chromosome 18 (left), and on SNPs from only within the focal region on chromosome 18 (right). SNPs were retained if they passed depth and quality filters and provided that at least 90% of individuals had genotypes with at least 3x coverage. Locality: Ko = Konstanz; Ra = Radolfzell; Ri = Rimbo; Up = Uppsala. Taxon: C.C. = carrion crow (*Corvus (corone) corone*); H.C. = hooded crow (*Corvus (corone) cornix*). Ordinations on SNPs analyzed with *SNPRelate* and visualized with *R* and the *tidyverse*.

**Fig S20.**
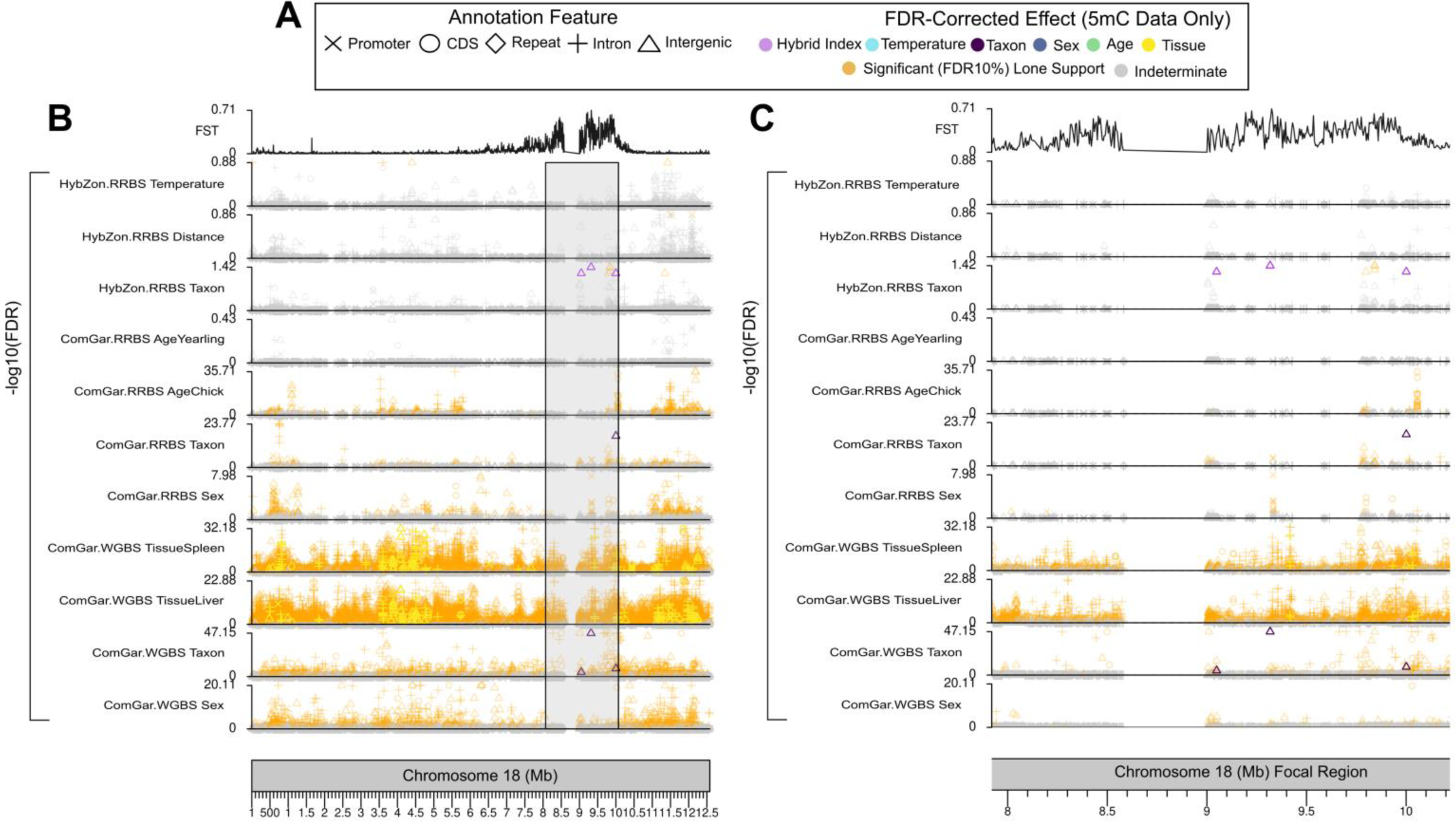
DMPs on chromosome 18 for all variables. DNA methylation (-log10(*p*-value)) and genetic differentiation (F_ST_) data for hooded, carrion, and hybrid crows, with a (A) common legend, showing chromosome 18 and the focal region. (B) CpGs from methylation experiments corresponding to the ComGar.WGBS, ComGar.RRBS, and HybZon.RRBS experiments along the bottom 11 panels showing the FDR-corrected p-value for each effect. Points are colored orange if they alone have an FDR < 10%, and grey if they are not significant for the target effect. Points are colored corresponding to the ultimate covariate classification if they have been validated multi-experimentally following the verified classification scheme (*cf.* Fig. 2). Top panel shows genetic differentiation between carrion and hooded crows. Visualized with *karyoploteR*.

**Fig S21.**
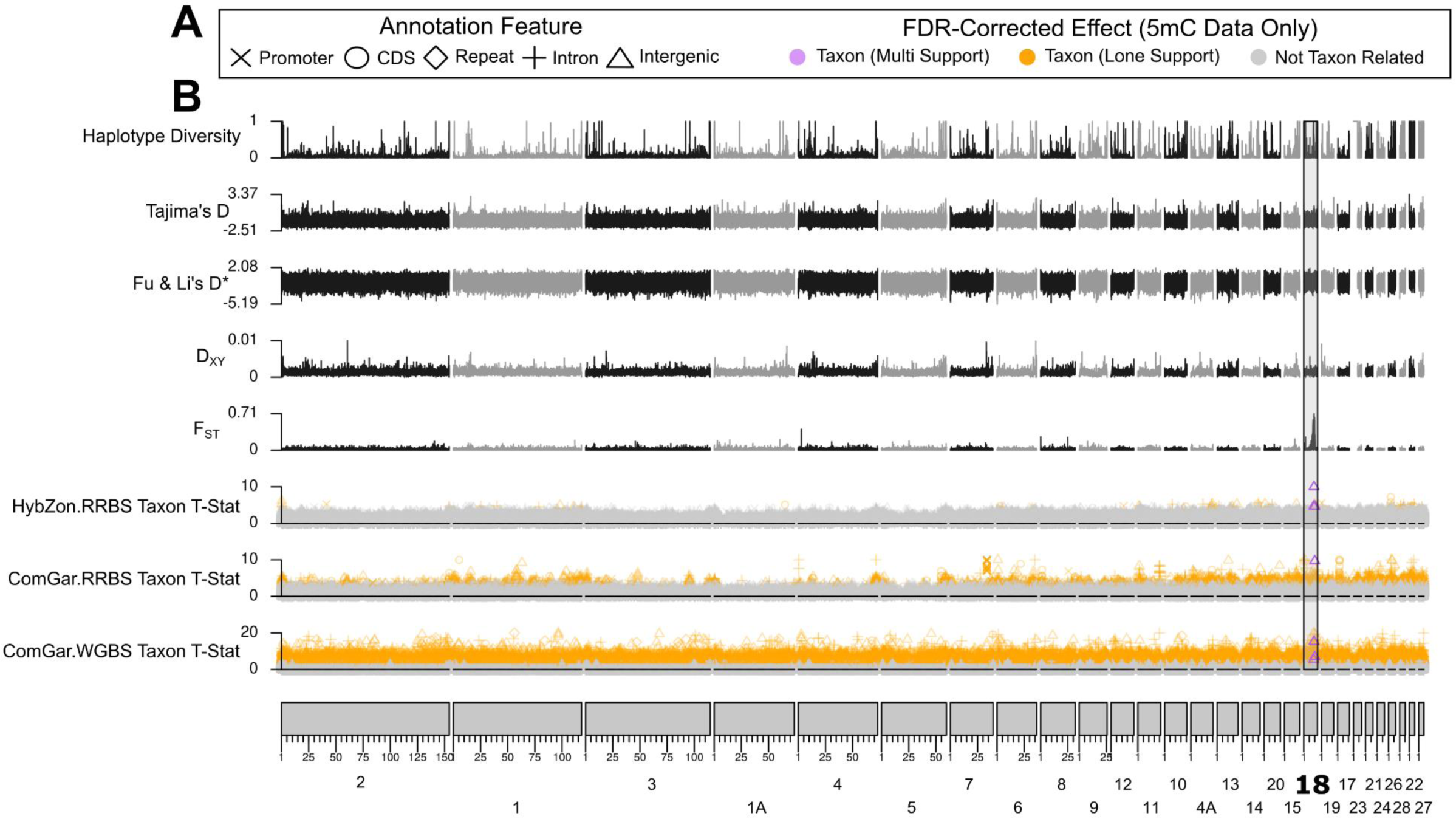
DMP test-statistics and genetic variation genome-wide. DNA methylation taxon-related test-statistics from DMP analyses and population genetic information for hooded, carrion, and hybrid crows genome-wide, with (**A**) a legend showing if the CpGs are supported by at least 2 experiments (multi-support), a single experiment, or no taxon effects (***cf.* Fig. 2**). CpGs were classified according to FDR-corrected *p-*values, divergence in means between groups, and inter-experimental validation (***cf.* Fig. 2**). (**B**) CpG-level DNA methylation test-statistics for taxon effects corresponding to the ComGar.WGBS, ComGar.RRBS, and HybZon.RRBS experiments along the bottom 3 panels. Points are colored orange if they alone have an FDR < 10%, and grey if they are not significant for the target effect. The top 5 panels show population genetic variation between (F_ST_, D_XY_) and among (Fu & Li’s D*, Tajima’s D, Haplotype Diversity) 28 whole-genome resequenced crows. Visualized with *karyoploteR*.

**Fig S22.**
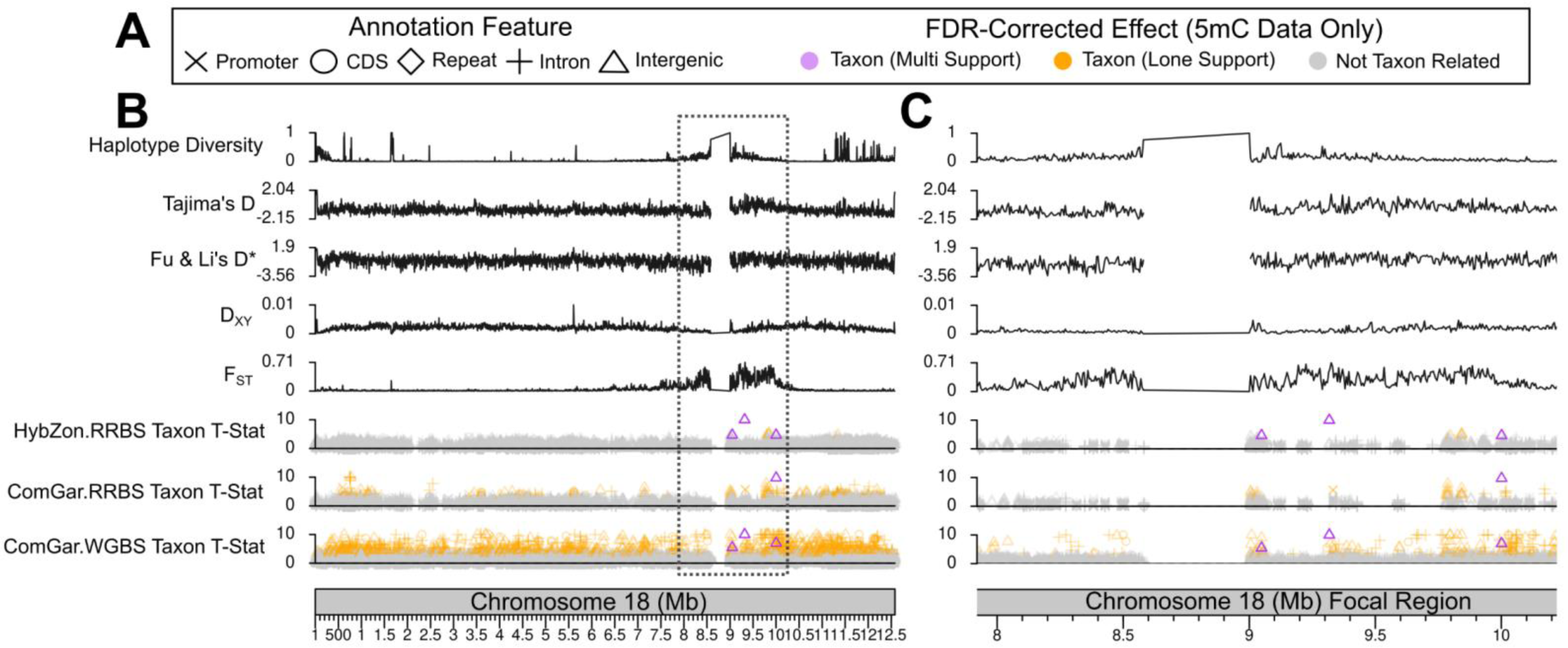
Test-statistics and genetic variation on chromosome 18. DNA methylation taxon-related test-statistics from DMP analyses and population genetic information for hooded, carrion, and hybrid crows across the (B) entire chromosome 18 and (C) within the focal region on chromosome 18, with (A) a legend showing if the CpGs are supported by at least 2 experiments (multi-support), a single experiment, or no taxon effects (*cf.* Fig. 2). CpGs were classified according to FDR-corrected *p-*values, divergence in means between groups, and inter-experimental validation (*cf.* Fig. 2). (B) CpG-level DNA methylation test-statistics for taxon effects corresponding to the ComGar.WGBS, ComGar.RRBS, and HybZon.RRBS experiments along the bottom 3 panels. Points are colored orange if they alone have an FDR < 10%, and grey if they are not significant for the target effect. The top 5 panels show population genetic variation between (F_ST_, D_XY_) and among (Fu & Li’s D*, Tajima’s D, Haplotype Diversity) 28 whole-genome resequenced crows. The portion with the putative centromere between 8.5 – 9-Mb has no information in any experiment (genetic data lines simply connect points on either side). These data were used as input for the supervised machine learning component to assess the chromosomal determinants on DNA methylation divergence (averaged in 5kb windows). Visualized with *karyoploteR*.

**Fig S23.**
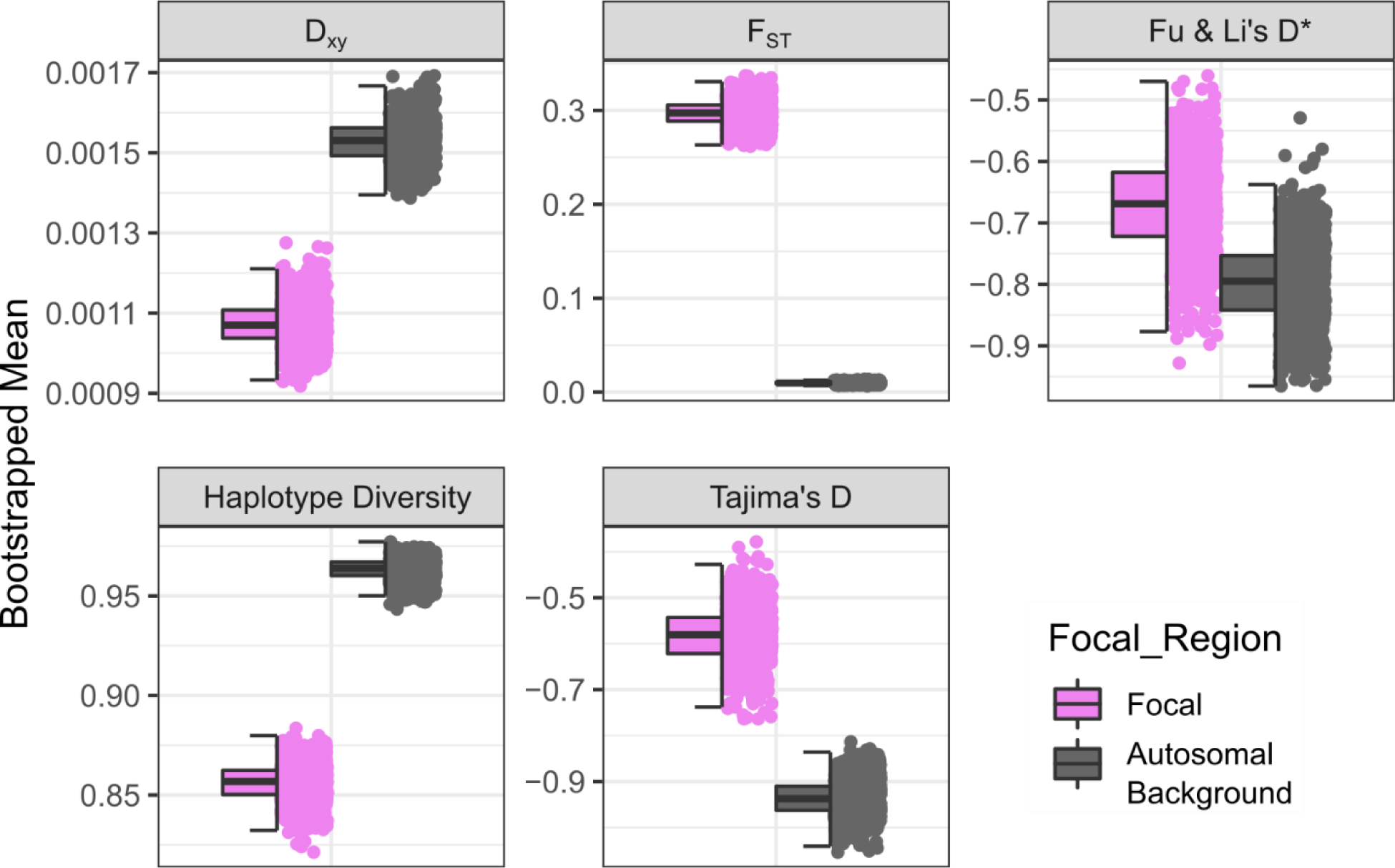
Genetic variation within and outside focal region. Population genetic variation, calculated in 5-Kb windows genome-wide from whole-genome resequencing data. D_XY_ and F_ST_ show divergence and differentiation between hooded and carrion crows, while the other metrics show overall variation across all crows. Distributions were created from 1,000 bootstrap sampling events of calculating the mean. Each bootstrap replicate sampled equal numbers of focal and autosomal background windows, which was equal to half the number of total available focal windows.

**Fig S24.**
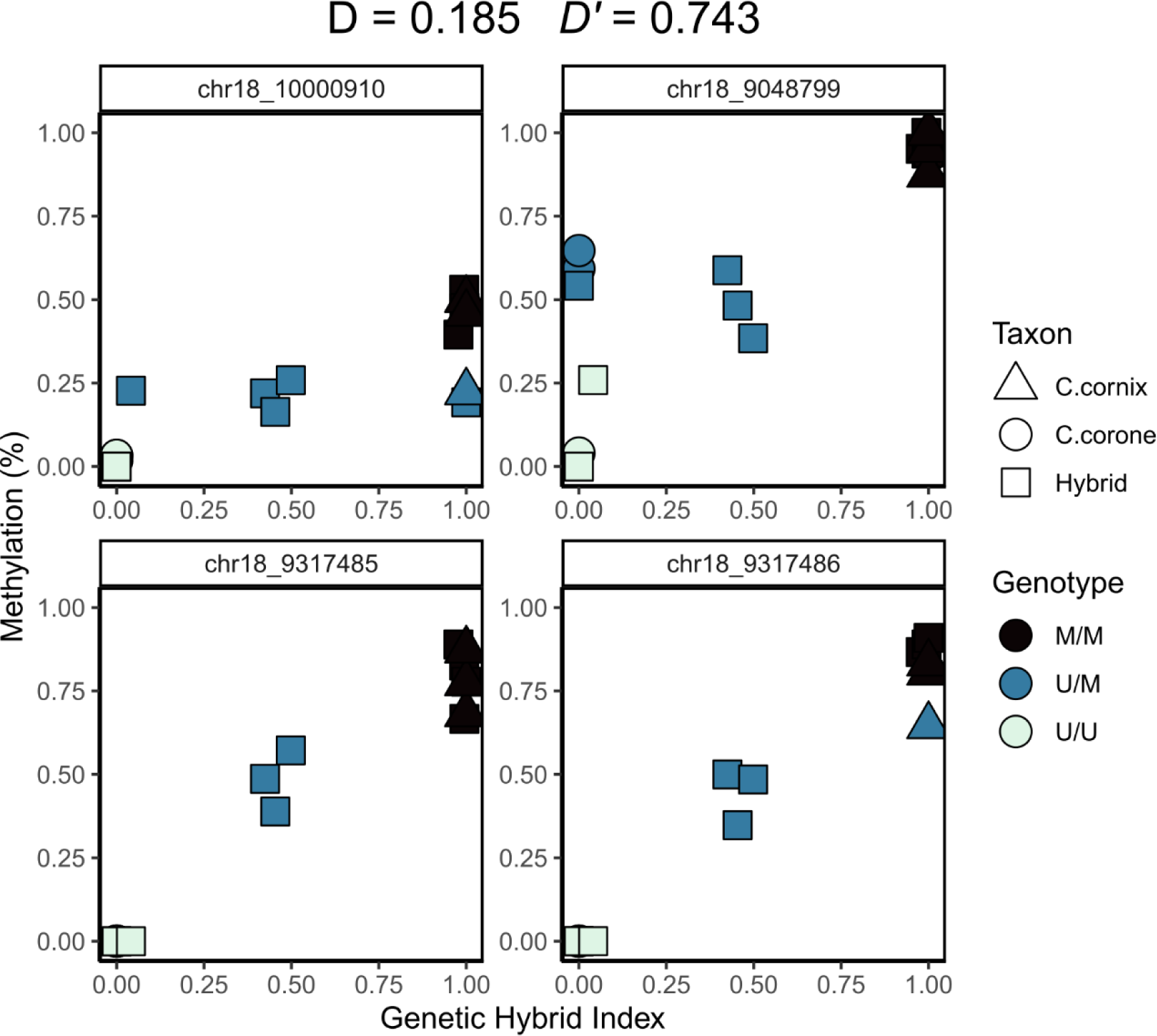
DNA methylation linkage disequilibrium at candidate taxon DMPs. Taxonomically relevant sites were identified from the multi-experimental classification approach, with the sites shown above (*n* = 4). For each site, each individual was assigned a methylation genotype based on k-means clustering of the methylation values (y-axis), resulting in the genotypes coloured above (epialleles M and U for methylated and unmethylated). LD between these sites was then calculated using classical genetics approaches, with all the analyses and plots done in *R* using the tidyverse.

## References

1. Wright S. Evolution in Mendelian Populations. Genetics. 1931;16: 97–159.

2. Jablonka E, Raz G. Transgenerational Epigenetic Inheritance: Prevalence, Mechanisms, and Implications for the Study of Heredity and Evolution. Q Rev Biol. 2009;84: 131–176. doi:10.1086/598822

3. Heard E, Martienssen RA. Transgenerational Epigenetic Inheritance: myths and mechanisms. Cell. 2014;157: 95–109. doi:10.1016/j.cell.2014.02.045

4. Fitz-James MH, Cavalli G. Molecular mechanisms of transgenerational epigenetic inheritance. Nat Rev Genet. 2022;23: 325–341. doi:10.1038/s41576-021-00438-5

5. Waddington CH. The Basic Ideas of Biology. Biol Theory. 2008;3: 238–253. doi:10.1162/biot.2008.3.3.238

6. Taudt A, Colomé-Tatché M, Johannes F. Genetic sources of population epigenomic variation. Nat Rev Genet. 2016;17: 319–332. doi:10.1038/nrg.2016.45

7. Christina L. Richards, Massimo Pigliucci. Epigenetic Inheritance. A Decade into the Extended Evolutionary Synthesis. Paradigmi. 2020; 463–494. doi:10.30460/99624

8. Zhang H, Lang Z, Zhu J-K. Dynamics and function of DNA methylation in plants. Nat Rev Mol Cell Biol. 2018;19: 489–506. doi:10.1038/s41580-018-0016-z

9. van der Graaf A, Wardenaar R, Neumann DA, Taudt A, Shaw RG, Jansen RC, et al. Rate, spectrum, and evolutionary dynamics of spontaneous epimutations. Proc Natl Acad Sci. 2015;112: 6676–6681. doi:10.1073/pnas.1424254112

10. Yao N, Schmitz RJ, Johannes F. Epimutations Define a Fast-Ticking Molecular Clock in Plants. Trends Genet. 2021;37: 699–710. doi:10.1016/j.tig.2021.04.010

11. Hazarika RR, Serra M, Zhang Z, Zhang Y, Schmitz RJ, Johannes F. Molecular properties of epimutation hotspots. Nat Plants. 2022;8: 146–156. doi:10.1038/s41477-021-01086-7

12. Kawakatsu T, Huang SC, Jupe F, Sasaki E, Schmitz RJ, Urich MA, et al. Epigenomic Diversity in a Global Collection of Arabidopsis thaliana Accessions. Cell. 2016;166: 492–505. doi:10.1016/j.cell.2016.06.044

13. Cubas P, Vincent C, Coen E. An epigenetic mutation responsible for natural variation in floral symmetry. Nature. 1999;401: 157–161. doi:10.1038/43657

14. Manning K, Tör M, Poole M, Hong Y, Thompson AJ, King GJ, et al. A naturally occurring epigenetic mutation in a gene encoding an SBP-box transcription factor inhibits tomato fruit ripening. Nat Genet. 2006;38: 948–952. doi:10.1038/ng1841

15. Furci L, Jain R, Stassen J, Berkowitz O, Whelan J, Roquis D, et al. Identification and characterisation of hypomethylated DNA loci controlling quantitative resistance in Arabidopsis. eLife. 2019;8: e40655. doi:10.7554/eLife.40655

16. Reik W, Dean W, Walter J. Epigenetic Reprogramming in Mammalian Development. Science. 2001;293: 1089–1093. doi:10.1126/science.1063443

17. Horsthemke B. A critical view on transgenerational epigenetic inheritance in humans. Nat Commun. 2018;9: 2973. doi:10.1038/s41467-018-05445-5

18. Beck D, Ben Maamar M, Skinner MK. Integration of sperm ncRNA-directed DNA methylation and DNA methylation-directed histone retention in epigenetic transgenerational inheritance. Epigenetics Chromatin. 2021;14: 6. doi:10.1186/s13072-020-00378-0

19. Leroux S, Gourichon D, Leterrier C, Labrune Y, Coustham V, Rivière S, et al. Embryonic environment and transgenerational effects in quail. Genet Sel Evol. 2017;49: 14. doi:10.1186/s12711-017-0292-7

20. Pierron F, Lorioux S, Héroin D, Daffe G, Etcheverria B, Cachot J, et al. Transgenerational epigenetic sex determination: Environment experienced by female fish affects offspring sex ratio. Environ Pollut. 2021;277: 116864. doi:10.1016/j.envpol.2021.116864

21. Thorson JLM, Beck D, Ben Maamar M, Nilsson EE, Skinner MK. Ancestral plastics exposure induces transgenerational disease-specific sperm epigenome-wide association biomarkers. Gerlinde M, editor. Environ Epigenetics. 2021;7: dvaa023. doi:10.1093/eep/dvaa023

22. Le Luyer J, Laporte M, Beacham TD, Kaukinen KH, Withler RE, Leong JS, et al. Parallel epigenetic modifications induced by hatchery rearing in a Pacific salmon. Proc Natl Acad Sci. 2017;114: 12964–12969. doi:10.1073/pnas.1711229114

23. Meröndun J, Murray DL, Shafer ABA. Genome-scale sampling suggests cryptic epigenetic structuring and insular divergence in Canada lynx. Mol Ecol. 2019; mec.15131. doi:10.1111/mec.15131

24. Wang Y, Qiao Z, Mao L, Li F, Liang X, An X, et al. Sympatric speciation of the spiny mouse from Evolution Canyon in Israel substantiated genomically and methylomically. Proc Natl Acad Sci. 2022;119: e2121822119. doi:10.1073/pnas.2121822119

25. Cossette M, Stewart DT, Haghani A, Zoller JA, Shafer ABA, Horvath S. Epigenetics and island-mainland divergence in an insectivorous small mammal. Mol Ecol. 2023;32: 152–166. doi:10.1111/mec.16735

26. Heckwolf MJ, Meyer BS, Häsler R, Höppner MP, Eizaguirre C, Reusch TBH. Two different epigenetic information channels in wild three-spined sticklebacks are involved in salinity adaptation. Sci Adv. 2020;6: eaaz1138. doi:10.1126/sciadv.aaz1138

27. Lindner M, Laine VN, Verhagen I, Viitaniemi HM, Visser ME, Oers K, et al. Rapid changes in DNA methylation associated with the initiation of reproduction in a small songbird. Mol Ecol. 2021;30: 3645–3659. doi:10.1111/mec.15803

28. Vernaz G, Hudson AG, Santos ME, Fischer B, Carruthers M, Shechonge AH, et al. Epigenetic divergence during early stages of speciation in an African crater lake cichlid fish. Nat Ecol Evol. 2022;6: 1940–1951. doi:10.1038/s41559-022-01894-w

29. Laporte M, Le Luyer J, Rougeux C, Dion-Côté A-M, Krick M, Bernatchez L. DNA methylation reprogramming, TE derepression, and postzygotic isolation of nascent animal species. Sci Adv. 2019;5: eaaw1644. doi:10.1126/sciadv.aaw1644

30. Taff CC, Campagna L, Vitousek MN. Genome-wide variation in DNA methylation is associated with stress resilience and plumage brightness in a wild bird. Mol Ecol. 2019;28: 3722–3737. doi:10.1111/mec.15186

31. Watson H, Powell D, Salmón P, Jacobs A, Isaksson C. Urbanization is associated with modifications in DNA methylation in a small passerine bird. Evol Appl. 2021;14: 85–98. doi:10.1111/eva.13160

32. Sepers B, Chen RS, Memelink M, Verhoeven KJF, Van Oers K. Variation in DNA Methylation in Avian Nestlings Is Largely Determined by Genetic Effects. Mulligan C, editor. Mol Biol Evol. 2023;40: msad086. doi:10.1093/molbev/msad086

33. Hanson HE, Wang C, Schrey AW, Liebl AL, Ravinet M, Jiang RHY, et al. Epigenetic Potential and DNA Methylation in an Ongoing House Sparrow (*Passer domesticus*) Range Expansion. Am Nat. 2022;200: 662–674. doi:10.1086/720950

34. Barton NH, Hewitt GM. Analysis of Hybrid Zones. Annu Rev Ecol Syst. 1985;16: 113–148. doi:10.1146/annurev.es.16.110185.000553

35. Gompert Z, Mandeville EG, Buerkle CA. Analysis of Population Genomic Data from Hybrid Zones. Annu Rev Ecol Evol Syst. 2017;48: 207–229. doi:10.1146/annurev-ecolsys-110316-022652

36. Knief U, Bossu CM, Saino N, Hansson B, Poelstra J, Vijay N, et al. Epistatic mutations under divergent selection govern phenotypic variation in the crow hybrid zone. Nat Ecol Evol. 2019;3: 570–576. doi:10.1038/s41559-019-0847-9

37. Baldassarre DT, White TA, Karubian J, Webster MS. Genomic and morphological analysis of a semipermeable avian hybrid zone suggests asymmetrical introgression of a sexual signal. Evolution. 2014;68: 2644–2657. doi:10.1111/evo.12457

38. Metzler D, Knief U, Peñalba JV, Wolf JBW. Assortative mating and epistatic mating-trait architecture induce complex movement of the crow hybrid zone. Evolution. 2021;75: 3154–3174. doi:10.1111/evo.14386

39. Poelstra JW, Vijay N, Bossu CM, Lantz H, Ryll B, Müller I, et al. The genomic landscape underlying phenotypic integrity in the face of gene flow in crows. Science. 2014;344: 1410–4. doi:10.1126/science.1253226

40. Vijay N, Bossu CM, Poelstra JW, Weissensteiner MH, Suh A, Kryukov AP, et al. Evolution of heterogeneous genome differentiation across multiple contact zones in a crow species complex. Nat Commun. 2016;7: 13195. doi:10.1038/ncomms13195

41. Weissensteiner MH, Pang AWC, Bunikis I, Höijer I, Vinnere-Petterson O, Suh A, et al. Combination of short-read, long-read, and optical mapping assemblies reveals large-scale tandem repeat arrays with population genetic implications. Genome Res. 2017;27: 697–708. doi:10.1101/gr.215095.116

42. Sepers B, van den Heuvel K, Lindner M, Viitaniemi H, Husby A, van Oers K. Avian ecological epigenetics: pitfalls and promises. J Ornithol. 2019 [cited 30 Aug 2019]. doi:10.1007/s10336-019-01684-5

43. Holtmann B, Buskas J, Steele M, Solokovskis K, Wolf JBW. Dominance relationships and coalitionary aggression against conspecifics in female carrion crows. Sci Rep. 2019;9: 1–8. doi:10.1038/s41598-019-52177-7

44. Blotzheim G von, Bauer KM, Bezzel E. Passeriformes (4. Teil): Corvidae – Sturnidae Rabenvögel, Starenvögel. AULA, Wiesbaden. 1993.

45. Griffith R, Double MC, Orr K, Dawson RJG. A DNA test to sex most birds. Mol Ecol. 1998;7: 1071–1075.

46. Gompert Z, Alex Buerkle C. introgress: a software package for mapping components of isolation in hybrids. Mol Ecol Resour. 2010;10: 378–384. doi:10.1111/j.1755-0998.2009.02733.x

47. Ioshikhes IP, Zhang MQ. Large-scale human promoter mapping using CpG islands. Nat Genet. 2000;26: 61–63. doi:10.1038/79189

48. Wu H, Caffo B, Jaffee HA, Irizarry RA, Feinberg AP. Redefining CpG islands using hidden Markov models. Biostatistics. 2010;11: 499–514. doi:10.1093/biostatistics/kxq005

49. Quinlan AR, Hall IM. BEDTools: a flexible suite of utilities for comparing genomic features. Bioinformatics. 2010;26: 841–842. 10.1093/bioinformatics/btq033

50. Smit A, Hubley R, Green P. RepeatMasker. 2013. Available: http://www.repeatmasker.org

51. R Core Team. R: A language and environment for statistical computing. R Foundation for Statistical Computing, Vienna, Austria. URL https://www.R-project.org/. 2017. p. Online.

52. Lepais O, Weir JT. SimRAD: An R package for simulation-based prediction of the number of loci expected in RADseq and similar genotyping by sequencing approaches. Mol Ecol Resour. 2014;14: 1314–1321. doi:10.1111/1755-0998.12273

53. Krueger F, Andrews SR. Bismark: A flexible aligner and methylation caller for Bisulfite-Seq applications. Bioinformatics. 2011;27: 1571–1572. doi:10.1093/bioinformatics/btr167

54. Barrow TM, Byun H-M. Single nucleotide polymorphisms on DNA methylation microarrays: precautions against confounding. Epigenomics. 2014;6: 577–579. doi:10.2217/epi.14.55

55. Bushnell B. BBTools: BBMap short read aligner, and other bioinformatic tools. 2021. Available: https://sourceforge.net/projects/bbmap/

56. Li H, Durbin R. Fast and accurate short read alignment with Burrows-Wheeler transform. Bioinformatics. 2009;25: 1754–1760. doi:10.1093/bioinformatics/btp324

57. Li H, Handsaker B, Wysoker A, Fennell T, Ruan J, Homer N, et al. The Sequence Alignment/Map format and SAMtools. Bioinformatics. 2009;25: 2078–2079. doi:10.1093/bioinformatics/btp352

58. Auwera GA, Carneiro MO, Hartl C, Poplin R, del Angel G, Levy-Moonshine A, et al. From FastQ Data to High-Confidence Variant Calls: The Genome Analysis Toolkit Best Practices Pipeline. Curr Protoc Bioinforma. 2013;43. doi:10.1002/0471250953.bi1110s43

59. Pedersen BS, Quinlan AR. Mosdepth: quick coverage calculation for genomes and exomes. Hancock J, editor. Bioinformatics. 2018;34: 867–868. doi:10.1093/bioinformatics/btx699

60. Danecek P, Bonfield JK, Liddle J, Marshall J, Ohan V, Pollard MO, et al. Twelve years of SAMtools and BCFtools. GigaScience. 2021;10: giab008. doi:10.1093/gigascience/giab008

61. Martin SH, Davey JW, Salazar C, Jiggins CD. Recombination rate variation shapes barriers to introgression across butterfly genomes. PLOS Biol. 2019;17: e2006288. doi:10.1371/journal.pbio.2006288

62. Pan Y, Liu P, Wang F, Wu P, Cheng F, Jin X, et al. Lineage-specific positive selection on *ACE2* contributes to the genetic susceptibility of COVID-19. Natl Sci Rev. 2022;9: nwac118. doi:10.1093/nsr/nwac118

63. Zheng X, Levine D, Shen J, Gogarten SM, Laurie C, Weir BS. A high-performance computing toolset for relatedness and principal component analysis of SNP data. Bioinformatics. 2012;28: 3326–3328. doi:10.1093/bioinformatics/bts606

64. Wickham H. ggplot2: Elegant Graphics for Data Analysis. Springer-Verlag New York; 2016. Available: https://ggplot2.tidyverse.org

65. Gel B, Serra E. karyoploteR: an R/Bioconductor package to plot customizable genomes displaying arbitrary data. Hancock J, editor. Bioinformatics. 2017;33: 3088–3090. doi:10.1093/bioinformatics/btx346

66. Krueger F. Trim Galore! : A wrapper tool around Cutadapt and FastQC to consistently apply quality and adapter trimming to FastQ files. Available at https://www.bioinformatics.babraham.ac.uk/projects/trim_galore/. 2012. p. Online.

67. Knief U, Bossu CM, Wolf JBW. Extra-pair paternity as a strategy to reduce the costs of heterospecific reproduction? Insights from the crow hybrid zone. J Evol Biol. 2020;33: 727–733. doi:10.1111/jeb.13607

68. Wickham H, Averick M, Bryan J, Chang W, McGowan L, François R, et al. Welcome to the Tidyverse. J Open Source Softw. 2019;4: 1686. doi:10.21105/joss.01686

69. Kuhn M, Wickham H. Tidymodels: a collection of packages for modeling and machine learning using tidyverse principles. 2020. Available: https://www.tidymodels.org

70. Anderson MJ, Willis TJ. Canonical analysis of principal coordinates: a useful method of constrained ordination for ecology. Ecology. 2003;84: 511–525. doi:10.1890/0012-9658(2003)084[0511:CAOPCA]2.0.CO;2

71. Park Y, Wu H. Differential methylation analysis for BS-seq data under general experimental design. Bioinformatics. 2016;32: 1446–1453. doi:10.1093/bioinformatics/btw026

72. Angeloni A, Fissette S, Kaya D, Hammond JM, Gamaarachchi H, Deveson IW, et al. Extensive DNA methylome rearrangement during early lamprey embryogenesis. Nat Commun. 2024;15: 1977. doi:10.1038/s41467-024-46085-2

73. Chen L, Cheng Y, Zhang G, Zhou Y, Zhang Z, Chen Q, et al. WGBS of embryonic gonads revealed that long non-coding RNAs in the MHM region might be involved in cell autonomous sex identity and female gonadal development in chickens. Epigenetics. 2024;19: 2283657. doi:10.1080/15592294.2023.2283657

74. Gore AV, Tomins KA, Iben J, Ma L, Castranova D, Davis AE, et al. An epigenetic mechanism for cavefish eye degeneration. Nat Ecol Evol. 2018;2: 1155–1160. doi:10.1038/s41559-018-0569-4

75. Laine VN, Gossmann TI, Schachtschneider KM, Garroway CJ, Madsen O, Verhoeven KJF, et al. Evolutionary signals of selection on cognition from the great tit genome and methylome. Nat Commun. 2016;7: 1–9. doi:10.1038/ncomms10474

76. De Carvalho CF, Slate J, Villoutreix R, Soria-Carrasco V, Riesch R, Feder JL, et al. DNA methylation differences between stick insect ecotypes. Mol Ecol. 2023;32: 6809–6823. doi:10.1111/mec.17165

77. El Kamouh M, Brionne A, Sayyari A, Lallias D, Labbé C, Laurent A. Strengths and limitations of reduced representation bisulfite sequencing (RRBS) in the perspective of DNA methylation analysis in fish: a case-study on rainbow trout spermatozoa. Fish Physiol Biochem. 2024 [cited 17 May 2024]. doi:10.1007/s10695-024-01326-5

78. Meyer BS, Moiron M, Caswara C, Chow W, Fedrigo O, Formenti G, et al. Sex-specific changes in autosomal methylation rate in ageing common terns. Front Ecol Evol. 2023;11: 982443. doi:10.3389/fevo.2023.982443

79. Capra E, Turri F, Lazzari B, Biffani S, Lange Consiglio A, Ajmone Marsan P, et al. CpG DNA methylation changes during epididymal sperm maturation in bulls. Epigenetics Chromatin. 2023;16: 20. doi:10.1186/s13072-023-00495-6

80. Lundregan SL, Mäkinen H, Buer A, Holand H, Jensen H, Husby A. Infection by a helminth parasite is associated with changes in DNA methylation in the house sparrow. Ecol Evol. 2022;12: e9539. doi:10.1002/ece3.9539

81. Metzger DCH, Schulte PM. Persistent and plastic effects of temperature on DNA methylation across the genome of threespine stickleback (*Gasterosteus aculeatus*). Proc R Soc B Biol Sci. 2017;284: 20171667. doi:10.1098/rspb.2017.1667

82. Wright MN, Ziegler A. ranger : A Fast Implementation of Random Forests for High Dimensional Data in *C++* and *R*. J Stat Softw. 2017;77. doi:10.18637/jss.v077.i01

83. Chen T, Guestrin C. XGBoost: A Scalable Tree Boosting System. Proceedings of the 22nd ACM SIGKDD International Conference on Knowledge Discovery and Data Mining. San Francisco California USA: ACM; 2016. pp. 785–794. doi:10.1145/2939672.2939785

84. Kuhn M. Building Predictive Models in *R* Using the caret Package. J Stat Softw. 2008;28. doi:10.18637/jss.v028.i05

85. Greenwell B M, Boehmke B C. Variable Importance Plots—An Introduction to the vip Package. R J. 2020;12: 343. doi:10.32614/RJ-2020-013

86. Altmann A, Toloşi L, Sander O, Lengauer T. Permutation importance: a corrected feature importance measure. Bioinformatics. 2010;26: 1340–1347. doi:10.1093/bioinformatics/btq134

87. Poelstra JW, Vijay N, Hoeppner MP, Wolf JBW. Transcriptomics of colour patterning and coloration shifts in crows. Mol Ecol. 2015;24: 4617–4628. doi:10.1111/mec.13353

88. Sheldon EL, Schrey Aaron W, Hurley LL, Griffith SC. Dynamic changes in DNA methylation during postnatal development in zebra finches *Taeniopygia guttata* exposed to different temperatures. J Avian Biol. 2020;51: jav.02294. doi:10.1111/jav.02294

89. Gama-Sosa MA, Midgett RM, Slagel VA, Githens S, Kuo KC, Gehrke CW, et al. Tissue-specific differences in DNA methylation in various mammals. Biochim Biophys Acta. 1983;740: 212–219. doi:10.1016/0167-4781(83)90079-9

90. Izzo F, Lee SC, Poran A, Chaligne R, Gaiti F, Gross B, et al. DNA methylation disruption reshapes the hematopoietic differentiation landscape. Nat Genet. 2020;52: 378–387. doi:10.1038/s41588-020-0595-4

91. De Paoli-Iseppi R, Deagle BE, Polanowski AM, McMahon CR, Dickinson JL, Hindell MA, et al. Age estimation in a long-lived seabird (*Ardenna tenuirostris*) using DNA methylation-based biomarkers. Mol Ecol Resour. 2019;19: 411–425. doi:10.1111/1755-0998.12981

92. Sun D, Layman TS, Jeong H, Chatterjee P, Grogan K, Merritt JR, et al. Genome-wide variation in DNA methylation linked to developmental stage and chromosomal suppression of recombination in white-throated sparrows. Mol Ecol. 2021;30: 3453–3467. doi:10.1111/mec.15793

93. Wu C-C, Klaesson A, Buskas J, Ranefall P, Mirzazadeh R, Söderberg O, et al. *In situ* quantification of individual mRNA transcripts in melanocytes discloses gene regulation of relevance to speciation. J Exp Biol. 2019;222: jeb.194431. doi:10.1242/jeb.194431

94. Bird A, Taggart M, Frommer M, Miller OJ, Macleod D. A fraction of the mouse genome that is derived from islands of nonmethylated, CpG-rich DNA. Cell. 1985;40: 91–99. doi:10.1016/0092-8674(85)90312-5

95. Long HK, King HW, Patient RK, Odom DT, Klose RJ. Protection of CpG islands from DNA methylation is DNA-encoded and evolutionarily conserved. Nucleic Acids Res. 2016;44: 6693–6706. doi:10.1093/nar/gkw258

96. Kawakami T, Smeds L, Backström N, Husby A, Qvarnström A, Mugal CF, et al. A high-density linkage map enables a second-generation collared flycatcher genome assembly and reveals the patterns of avian recombination rate variation and chromosomal evolution. Mol Ecol. 2014;23: 4035–4058. doi:10.1111/mec.12810

97. Carja O, MacIsaac JL, Mah SM, Henn BM, Kobor MS, Feldman MW, et al. Worldwide patterns of human epigenetic variation. Nat Ecol Evol. 2017;1: 1577–1583. doi:10.1038/s41559-017-0299-z

98. Shirai K, Sato MP, Nishi R, Seki M, Suzuki Y, Hanada K. Positive selective sweeps of epigenetic mutations regulating specialized metabolites in plants. Genome Res. 2021;31: 1060–1068. doi:10.1101/gr.271726.120

99. Vilgalys T, Rogers J, Jolly C, Mukherjee S, Tung J. Evolution of DNA methylation in Papio baboons. Mulligan C, editor. Mol Biol Evol. 2018. doi:10.1093/molbev/msy227

100. Hawe JS, Wilson R, Schmid KT, Zhou L, Lakshmanan LN, Lehne BC, et al. Genetic variation influencing DNA methylation provides insights into molecular mechanisms regulating genomic function. Nat Genet. 2022;54: 18–29. doi:10.1038/s41588-021-00969-x

101. Yagound B, Smith NMA, Buchmann G, Oldroyd BP, Remnant EJ. Unique DNA Methylation Profiles Are Associated with cis-Variation in Honey Bees. Costantini M, editor. Genome Biol Evol. 2019;11: 2517–2530. doi:10.1093/gbe/evz177

102. Min JL, Hemani G, Hannon E, Dekkers KF, Castillo-Fernandez J, Luijk R, et al. Genomic and phenotypic insights from an atlas of genetic effects on DNA methylation. Nat Genet. 2021;53: 1311–1321. doi:10.1038/s41588-021-00923-x

103. Höglund A, Henriksen R, Fogelholm J, Churcher AM, Guerrero-Bosagna CM, Martinez-Barrio A, et al. The methylation landscape and its role in domestication and gene regulation in the chicken. Nat Ecol Evol. 2020;4: 1713–1724. doi:10.1038/s41559-020-01310-1

104. Cooper DN, Krawczak M. Cytosine methylation and the fate of CpG dinucleotides in vertebrate genomes. Hum Genet. 1989;83: 181–188. doi:10.1007/BF00286715

105. Zhou W, Liang G, Molloy PL, Jones PA. DNA methylation enables transposable element-driven genome expansion. Proc Natl Acad Sci. 2020;117: 19359–19366. doi:10.1073/pnas.1921719117

106. Feinberg AP, Irizarry RA. Stochastic epigenetic variation as a driving force of development, evolutionary adaptation, and disease. Proc Natl Acad Sci. 2010;107: 1757–1764. doi:10.1073/pnas.0906183107

107. Warmuth VM, Weissensteiner MH, Wolf JBW. Accumulation and ineffective silencing of transposable elements on an avian W Chromosome. Genome Res. 2022;32: 671–681. doi:10.1101/gr.275465.121

108. Uller T. Evolutionary perspectives on transgenerational epigenetics. Transgenerational Epigenetics. Elsevier; 2019. pp. 333–350. doi:10.1016/B978-0-12-816363-4.00015-8

109. Shafer ABA, Wolf JBW. Widespread evidence for incipient ecological speciation: a meta-analysis of isolation-by-ecology. Ecol Lett. 2013;16: 940–950. doi:10.1111/ele.12120

110. Corbett RJ, Te Pas MFW, Van Den Brand H, Groenen MAM, Crooijmans RPMA, Ernst CW, et al. Genome-Wide Assessment of DNA Methylation in Chicken Cardiac Tissue Exposed to Different Incubation Temperatures and CO2 Levels. Front Genet. 2020;11: 558189. doi:10.3389/fgene.2020.558189

111. Yan X, Liu H, Liu J, Zhang R, Wang G, Li Q, et al. Evidence in duck for supporting alteration of incubation temperature may have influence on methylation of genomic DNA. Poult Sci. 2015;94: 2537–2545. doi:10.3382/ps/pev201

112. Viitaniemi HM, Verhagen I, Visser ME, Honkela A, van Oers K, Husby A. Seasonal Variation in Genome-Wide DNA Methylation Patterns and the Onset of Seasonal Timing of Reproduction in Great Tits. Meyer M, editor. Genome Biol Evol. 2019;11: 970–983. doi:10.1093/gbe/evz044

113. Kheirkhah Rahimabad P, Arshad SH, Holloway JW, Mukherjee N, Hedman A, Gruzieva O, et al. Association of Maternal DNA Methylation and Offspring Birthweight. Reprod Sci. 2021;28: 218–227. doi:10.1007/s43032-020-00281-9

114. Pu N, Yang Q, Shi X-L, Chen W-W, Li X-Y, Zhang G-F, et al. Gene–environment interaction between APOA5 c.553G>T and pregnancy in hypertriglyceridemia-induced acute pancreatitis. J Clin Lipidol. 2020;14: 498–506. doi:10.1016/j.jacl.2020.05.003

115. Caizergues AE, Le Luyer J, Grégoire A, Szulkin M, Senar J, Charmantier A, et al. Epigenetics and the city: Non-parallel DNA methylation modifications across pairs of urban-forest Great tit populations. Evol Appl. 2022;15: 149–165. doi:10.1111/eva.13334

116. Morgan HD, Sutherland HGE, Martin DIK, Whitelaw E. Epigenetic inheritance at the agouti locus in the mouse. Nat Genet. 1999;23: 314–318. doi:10.1038/15490

